# Human neurons from Christianson syndrome iPSCs reveal allele-specific responses to rescue strategies

**DOI:** 10.1101/444232

**Authors:** Sofia B. Lizarraga, Abbie M. Maguire, Li Ma, Laura I. van Dyck, Qing Wu, Dipal Nagda, Liane L. Livi, Matthew F. Pescosolido, Michael Schmidt, Shanique Alabi, Mara H. Cowen, Paul Brito-Vargas, Diane Hoffman-Kim, Ece D. Gamsiz Uzun, Avner Schlessinger, Richard N. Jones, Eric M. Morrow

**Affiliations:** Department of Biological Sciences, University of South Carolina, Columbia, South Carolina, USA; Center for Childhood Neurotherapeutics, University of South Carolina, Columbia, South Carolina, USA; Department of Molecular Biology, Cell Biology and Biochemistry, Brown University, Providence, Rhode Island, USA; Hassenfeld Child Health Innovation Institute, Brown University, Providence, Rhode Island, USA; Department of Molecular Pharmacology, Physiology, and Biotechnology, Brown University, Providence, Rhode Island, USA; Department of Pharmacological Sciences, Icahn School of Medicine at Mount Sinai, New York, New York, USA; Carney Institute for Brain Science, Brown University, Providence, Rhode Island, USA; Center for Biomedical Engineering, Brown University, Providence, Rhode Island, USA; Department of Pathology and Laboratory Medicine, Warren Alpert Medical School of Brown University, Providence, Rhode Island, USA; Center for Computational Molecular Biology, Brown University, Providence, Rhode Island, USA; Department of Psychiatry and Human Behavior, Warren Alpert Medical School of Brown University, Providence, Rhode Island, USA; Developmental Disorders Genetics Research Program, Emma Pendleton Bradley Hospital, East Providence, Rhode Island, USA

## Abstract

Human genetic disorders provide a powerful lens to understanding the human brain. Induced pluripotent stem cells (iPSC) represent an important, new resource for mechanistic studies and therapeutic development. Christianson syndrome (CS), an X-linked neurological disorder with attenuation of brain growth postnatally (postnatal microcephaly), is caused by mutations in *SLC9A6,* the gene encoding endosomal Na^+^/H^+^ exchanger 6 (NHE6). We developed CS iPSC lines from patients with a mutational spectrum, as well as robust biologically-related and isogenic controls. We demonstrate that mutations in CS lead to loss of protein function by a variety of mechanisms. Regardless of mutation, all patient-derived neurons demonstrate reduced neurite growth and arborization, likely underlying diminished postnatal brain growth in patients. Additionally, phenotype rescue strategies show allele-specific responses: a gene replacement strategy shows efficacy in nonsense mutations but not in a missense mutation, whereas application of exogenous trophic factors (BDNF or IGF-1) rescues arborization phenotypes across all mutations. Our data emphasize the important principle of personalized medicine whereby success of some therapeutic strategies may be more linked to patient genotype than others.

---

Christianson syndrome (CS) is an X-linked neurological disorder characterized by impaired cognitive development and epilepsy, with a distinctive social communication phenotype^1,2^. CS is caused by diverse mutations in *SLC9A6,* the gene encoding the endosomal Na^+^/H^+^ exchanger 6 (NHE6)^2,3^. Given the social phenotype, which includes a happy demeanor and autistic features, and the presentation of seizures and postnatal microcephaly, CS was originally termed X-linked Angelman syndrome (AS)^3^. Postnatal microcephaly (as opposed to primary microcephaly) is diagnosed when children are born with typical head circumference and subsequent attenuation of the expected brain growth in the first years of life^4^. The human brain increases in size by approximately two- to three-fold in the first years of life^5-7^, coincident with the emergence of experience-dependent acquisition of brain function and behavior. Thereby, study of human genetic disorders with postnatal microcephaly, such as CS or AS, may allow for the identification of genes involved in early childhood brain development, including genes that govern cognitive, motor, and social development^4^. As neurogenesis is largely complete in human brain prior to birth, postnatal microcephaly genes often regulate late neurodevelopmental processes involved in neuronal connectivity such as neuronal axonogenesis, axonal and dendritic arborization, and synaptic development.

Endosomes play an important role in the development of neuronal connectivity via a range of mechanisms, including by their involvement in the trafficking, recycling, and degradation of cargo, as well as by serving in endosomal signaling^8,9^. In particular, neurotrophin signaling, such as through the brain-derived neurotrophic factor (BDNF)/tropomyosin-related kinase B (TrkB) pathway, represents the classically studied endosomal signaling mechanism essential for neurite outgrowth and arborization^10^. Tight regulation of intra-endosomal proton concentration serves an essential role in endo-lysosomal maturation and function^11,12^. The vacuolar H+-ATPase (V-ATPase) is a pump that mediates acidification of endosomes and lysosomes^13^. The endosomal NHEs (particularly NHE6 and NHE9) allow proton efflux, and counter the V-ATPase by regulating relative alkalization of the lumen^14^. We and others have identified a role for intra-endosomal pH in modulation of neurotrophin signaling that drives neuronal arborization^15,16^. Loss of NHE6 in primary mouse neurons revealed a unique endosomal phenotype, namely, over-acidification of the endosome lumen^15^. Reduced endosomal pH in developing NHE6-null neurons was associated with attenuation of BDNF signaling via TrkB and diminished neuronal arborization in vitro and in vivo. Importantly, exogenous addition of BDNF to NHE6-null murine cultures rescued the arborization defect, consistent with the previously shown pH-dependent decrease in ligand-receptor binding^16^.

We and others have also recently shown that, in addition to neurodevelopmental disabilities, CS is associated with a progressive neurodegeneration phenotype^2,17,18^. Given the disease burden experienced by people with CS, we sought to develop an induced pluripotent stem cell (iPSC) system that could be utilized to study the cellular mechanisms of CS in human neurons, as well as serve for therapeutic development. Patient-derived iPSCs offer the opportunity to dissect mechanisms mediated by specific human mutations from the endogenous genomic locus. As has been well demonstrated recently for therapeutic development for cystic fibrosis, specific patient mutations show evidence of allele-specific responses to treatments^19,20^. Given our prior data on BDNF, one rescue strategy of interest is the exogenous (i.e., cell-non-autonomous) addition of trophic factors. Such factors might include not only BDNF but also insulin-like growth factor-1 (IGF-1). BDNF and IGF-1 have demonstrated activity on neurite growth and arborization^21^, and further still, IGF-1 is in clinical trials for other neurodevelopmental disorders with postnatal microcephaly and/or autistic features (e.g., NCT01970345, NCT01525901; clinicaltrials.gov). We also examine a rescue strategy involving cell-autonomous replacement of NHE6 via cDNA transfer. This therapeutic strategy may be pursued in theory through gene replacement or perhaps through upregulation of the related endosomal NHE9 in CS patients.

In this study, we developed a substantial iPSC resource to study CS disease mechanisms and therapeutic responses. This resource includes an allelic series of patient mutations and both genetically related and isogenic controls. Here, we utilize this resource to determine allele-specific action of mutations in patient-derived neurons, as well as distinct allele-specific responses to rescue strategies. While exogenous addition of BDNF or IGF-1 to patient neurons rescues neuronal phenotypes regardless of patient mutation, gene replacement strategies show successful rescue in a fashion that is dependent on the nature of the patient mutation.

## Results

### A majority of CS mutations lead to loss of protein through Nonsense mediated decay mechanisms

In order to establish a patient-derived cellular model for translational research in CS, we reprogrammed peripheral blood mononuclear cells (PBMCs) from five families, each of which contained a distinct mutation in *SLC9A6* in the affected boy, into iPSCs^22,23^ (Table 1). A majority of NHE6 mutations in CS appear to be loss-of-function (i.e., putatively protein truncating due to early frameshift, nonsense, or splicing mutations), while several missense or in-frame deletions have also been reported^2^. Our iPSC collection includes an allelic series of four distinct truncating mutations and one missense mutation in NHE6, the latter of which we hypothesize could also affect mRNA splicing. These mutations are as follows: Family 1, a single base pair duplication in exon 11 leading to a frameshift and premature stop codon between predicted transmembrane domains (TM) 10 and 11 (c.1414dupA, p.R472fsX4); Family 2, a nonsense mutation in exon 12 in predicted TM12 (c.1568G>A, p.W523X); Family 3, a missense mutation at the first base pair of exon 9 in predicted TM8 (c.1148G>A, p.G383D); Family 4, an eight base pair duplication in exon 3 leading to a frameshift and premature stop codon in predicted TM4 (c.540_547dupAGAAGTAT, p.F183fsX1); and Family 5, a nonsense mutation in exon 14 located in the cytoplasmic tail (c.1710G>A, p.W570X)^2^ (Fig. 1a; Table 1; and **Supplementary Figs. 1 and 2**). iPSCs were also derived from each proband’s genetically related, non-carrier male sibling for use as a paired control. Control and CS iPSCs had similar morphologies to human embryonic stem cells (hESCs), expressed endogenous pluripotency markers, had normal karyotypes, formed embryoid bodies *in vitro*, and formed teratomas *in vivo*, thereby demonstrating successful reprogramming (**Supplementary Table 1** and **Supplementary Figs. 3-10**).

**Figure 1.**
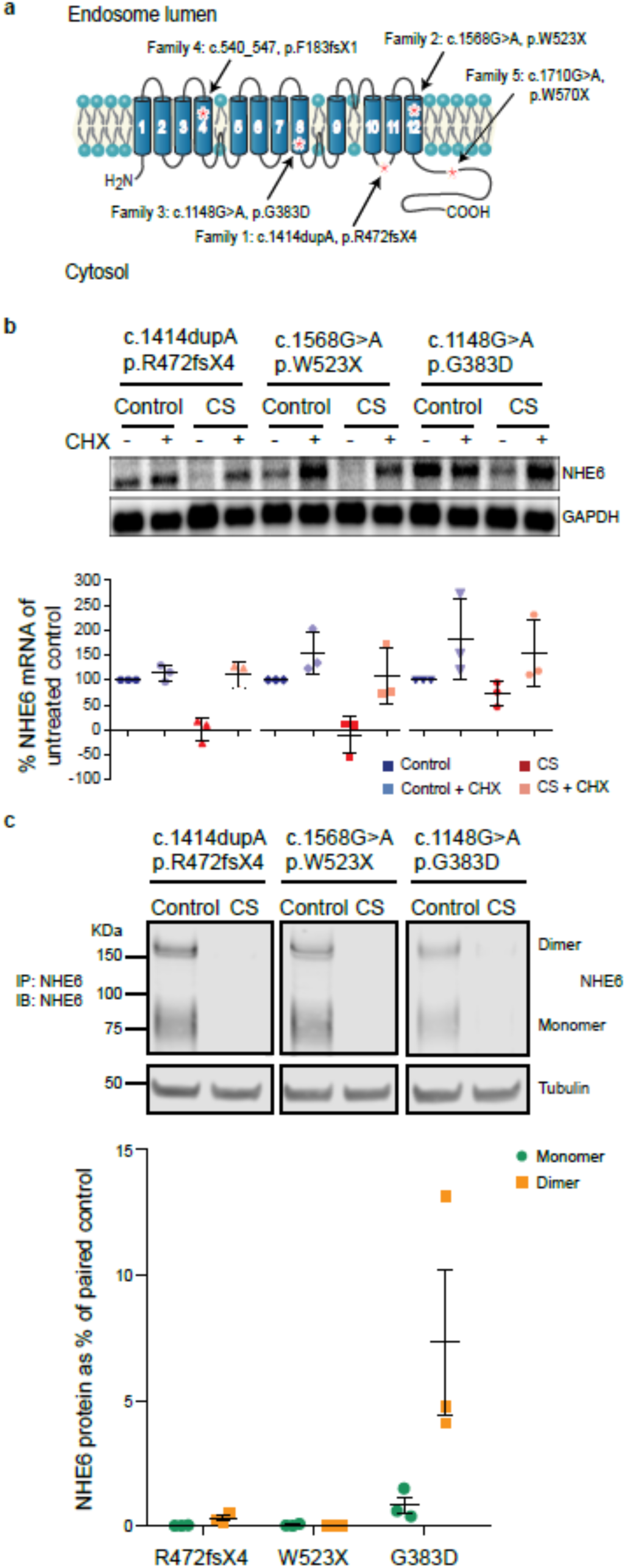
Establishment of iPSCs from CS patients with NHE6 mutations and from related controls. (**a**) iPSCs were derived from five CS patients and from their genetically related, unaffected brothers without NHE6 mutation as controls. The locations of the proband mutations are based on NHE6 transcript NM_001042537. NHE6 TM were predicted by the transmembrane hidden Markov model (TMHMM)^57^ for Ensembl transcript ENST00000370695 and protein ENSP00000359729 ^58^. (**b**) NHE6 mRNA quantification in CS iPSCs. Northern blotting was performed using samples from CS and control iPSCs treated (+) or untreated (-) with CHX. Expression of *SLC9A6* mRNA was normalized to *GAPDH* and plotted as a percentage of untreated control cells. *n* = 3 experiments using iPSCs of subsequent passages, mean ± s.e.m. (**c**) NHE6 protein quantification in CS iPSCs. In order to generate a relatively pure sample of endogenously expressed NHE6, CS and control iPSCs were immunoprecipitated for NHE6 followed by western blot analysis for NHE6. Expression of NHE6 dimer and monomer is plotted as a percentage of the paired control. Note that the anti-NHE6 antibody is a custom-made polyclonal sera to a cytoplasmic tail epitope (see Methods). Tubulin was used as a loading control. *n* = 3 experiments using iPSCs of subsequent passages, mean ± s.e.m.

**Table 1.**
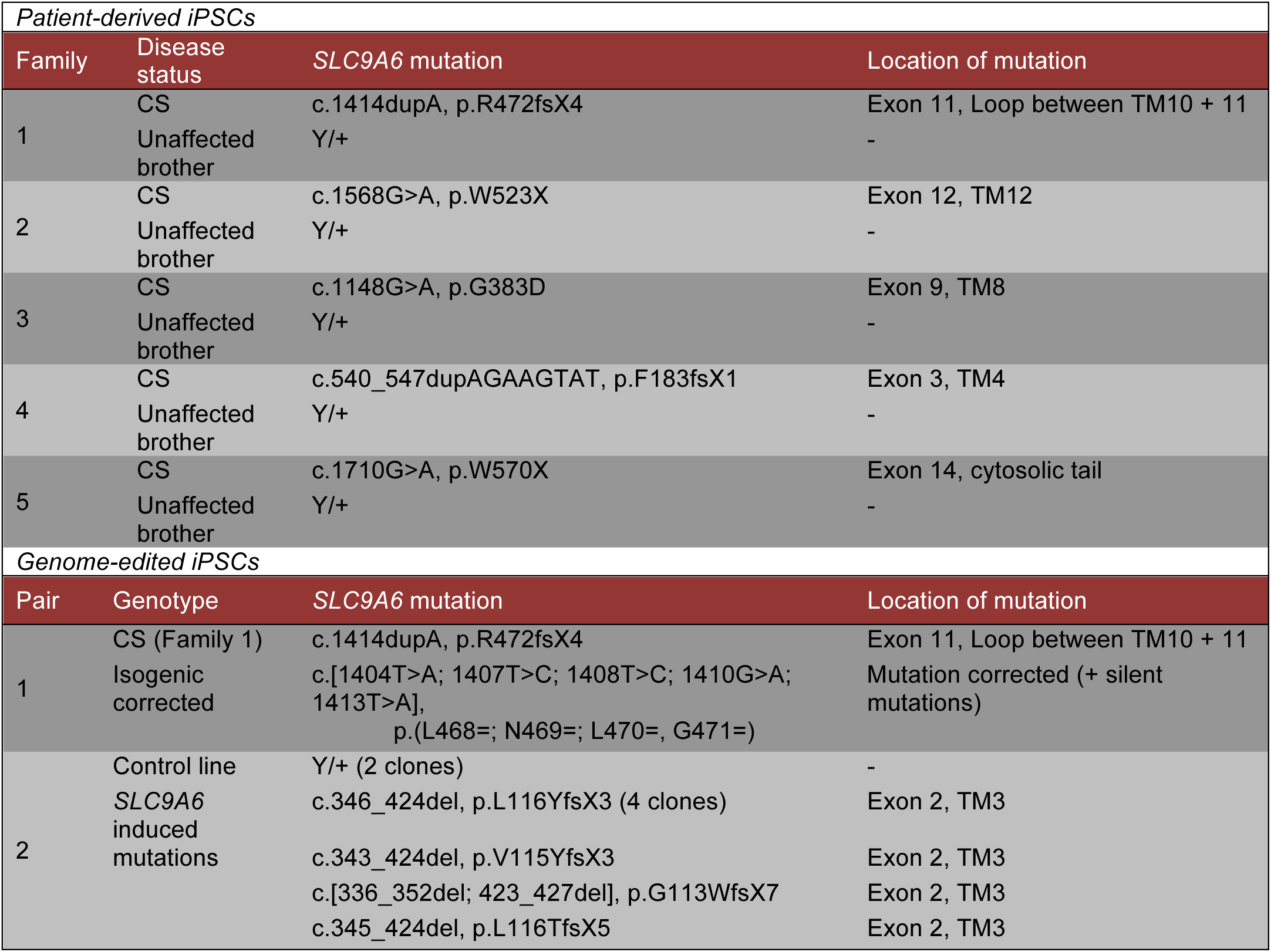
iPSC resources for study of CS. Resources include iPSCs derived from: 1) five families with distinct mutations, including paired CS patient and biologically related, non-carrier, unaffected male siblings; 2) isogenic CS and genome-corrected lines; and 3) isogenic control and *SLC9A6* induced-mutation lines. Mutations in *SLC9A6* are listed based on Ensembl transcript ENST00000370695. Transmembrane domains (TM) were predicted using TMHMM based on ENSP00000359729.

To determine the effects of the patient mutations on NHE6 gene and protein expression, we conducted northern and western blot analyses of representative mutations and subclonal lines. Notably, in all four of the putative protein-truncating mutations, we found *SLC9A6* mRNA to be absent. For example, in the CS patients mutations c.1414dupA/p.R472fsX4 and c.1568G>A/p.W523X, negligible amounts of mRNA were detectable by northern blot (Fig. 1b). To test if *SLC9A6* mRNA is degraded by nonsense-mediated decay (NMD), we treated iPSCs with cycloheximide (CHX), which prevents NMD by inhibiting translation and thereby interfering with recognition of a premature stop codon. Upon CHX treatment, the *SLC9A6* transcript was recovered in CS lines with these protein-truncating mutations (Fig. 1b). This result was corroborated at the level of protein analysis. Using western blot on NHE6 immunoprecipitates, NHE6 protein appeared to be absent in all CS lines carrying gene nonsense/frameshift mutations, for example, as shown for the c.1414dupA/p.R472fsX4 (Family 1) and c.1568G>A/p.W523X (Family 2) mutations (Fig. 1c). Similar northern blot and western blot results were obtained for the CS lines carrying the frameshift mutation c.540_547dupAGAAGTAT/p.F183fsX1 (Family 4) or the late nonsense mutation c.1710G>A/p.W570X (Family 5) (**Supplementary Fig. 11**). Therefore, these results establish that the majority of CS mutations, namely those leading to nonsense mutations, are likely to cause disease by a loss-of-function mechanism and that truncated NHE6 proteins are unlikely to be translated in large quantities.

### Missense G383D mutation demonstrates complex effects on mRNA and protein products

Interestingly, the NHE6 gene and protein in patient cells with the missense mutation c.1148G>A/p.G383D behaved differently than the nonsense mutations. The NHE6 mRNA transcript in cells with the missense mutation was not absent but instead was reduced to 58% ± 18.4% of control levels; however, the transcript level was recoverable with CHX treatment, suggesting some component of NMD (Fig. 1b, four right lanes on northern blot). Very low levels of protein were detectable in the mutant: 0.82% ± 0.34% of NHE6 monomer and 7.33% ± 2.90% of NHE6 dimer remained in CS lines with the c.1148G>A/p.G383D mutation as compared to control (Fig. 1c, right-most panel of western blot). Near complete lack of the monomer suggests that it may be unstable, whereas the dimer may be relatively more stable, although the dimer is also substantially decreased. Notably, in addition to creating a missense coding change, the c.1148G>A/p.G383D mutation occurs at the first base pair of exon 9 and therefore directly adjacent to the 3’ splice acceptor site (**Supplementary Fig. 1**). We predicted that this mutation may disrupt splicing, namely, skipping of exon 9 and splicing of exon 8 to exon 10, thereby leading to a frameshift and premature stop codon two codons downstream of the exon 8/10 junction (Fig. 2a). RT-PCR of cDNA from iPSCs with the c.1148G>A/p.G383D mutation, including sequencing of the PCR product, revealed the presence of an alternative splicing event generated by skipping exon 9 (Fig. 2a). Therefore, two mRNAs are produced, one with a nonsense mutation (subject to NMD and recoverable by cycloheximide), as well as an mRNA which will encode a protein product with the G383D missense mutation.

**Figure 2.**
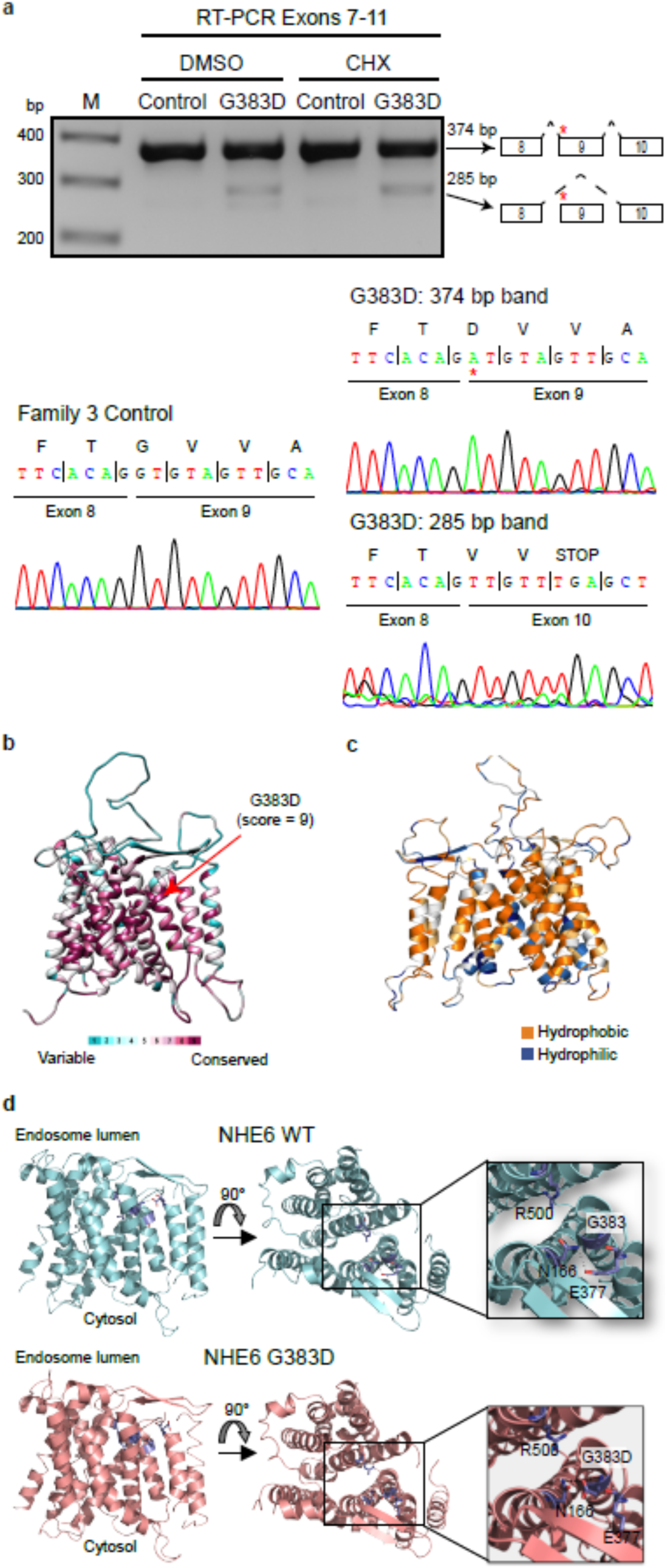
Analysis of complex alternative splicing, missense mutation. (**a**) The c.1148G>A/G383D mutation generates an alternative transcript by exon skipping. RT-PCR was performed on cDNA collected from iPSCs treated or untreated with CHX. The resulting bands were sequenced and revealed the presence of an alternative splicing event in the c.1148G>A/G383D mutation in which exon 9 is skipped. (**b**) Conservation of NHE6 by ConSurf analysis. Variable and conserved regions (turquoise through maroon) are mapped onto the NHE6 structural model. (**c**) Hydrophobicity analysis of the NHE6 structural model. Hydrophobic (orange) residues are located in TM, while hydrophilic (blue) residues are located in loop regions. (**d**) Structural modeling of the c.1148G>A/G383D mutation. The model of NHE6 predicts that the c.1148G>A/p.G383D mutation disrupts helix packing between TM8 and TM2, possibly due to hydrogen bonding of D383 with N166 and loss of an interaction between N166 and E377.

With regard to the G383D mutation, through use of structural modeling of the protein, we investigated if this missense mutation could affect protein structure. We constructed a homology model of NHE6 based on the crystal structure of the *Escherichia coli* Na^+^/H^+^ exchanger NhaA^24^, which was previously used to model other members of the SLC9 family^25^. Consurf analysis, which calculates the rate of evolution at each amino acid based on phylogenetic relations between homologous sequences, shows the highest conservation score for the G383 residue (Fig. 2b); this hints at the importance of this residue for the proper functioning of NHE6. For example, G383 is highly conserved across human, yeast, and bacterial NHE homologs (**Supplementary Fig. 12**). The NHE6 structural model shows that, similar to other membrane proteins, hydrophobic residues are located in the transmembrane domains (TM), while polar residues are located in loops (Fig. 2c); these results increase our confidence in the model. The model predicts that the G383 residue is positioned toward the endosomal end of NHE6 facing the core of the transporter on TM8. This domain has been proposed to function in the regulation of ion transport mediated by NHE family members^26^. Mutation of this non-polar glycine residue to an acidic aspartate residue is therefore likely to impact the transporter’s structure by disrupting helix packing between TM7 and TM8. Interestingly, modeling of the G383D variant suggests that this residue may also disrupt the transporter structure by forming a hydrogen bond with N166 on TM2 (Fig. 2d). Alternatively, G383D variant could form a salt-bridge with R500 on TM11 (not shown), thereby stabilizing the inward open NHE6 conformation. Taken together, structural modeling suggests that the G383D mutation hampers the structure and dynamics of TMs of NHE6, negatively affecting ion transport. In summary, the G383D mutation is predicted to be loss-of-function by complex mechanisms including NMD and defects in splicing and protein stability, resulting in a near complete loss of protein. The small amount of protein remaining is predicted to have impaired transport based on structural analyses. However, it will be important to address the possibility of a dominant-negative action of the remnant protein, as this will have important significance to therapies involving gene replacement strategies.

### Generation of isogenic control lines using CRISPR/Cas9-based genome editing

In order to generate a comprehensive resource for the study of disease-relevant NHE6 mutations in human iPSCs, we also generated pairs of isogenic lines (mutation and control) through genome editing using a Clustered Regularly Interspaced Short Palindromic Repeats (CRISPR)/Cas9 technique. We first successfully generated a line wherein the c.1414dupA/p.R472fsX4 mutation was corrected back to reference sequence (Table 1 and **Supplementary Fig. 13a**). In order to efficiently generate corrected clones produced by homology-directed repair, silent mutations were introduced into the guide RNA to prevent re-editing of the target sequence^27^. Immunoprecipitation of NHE6 followed by western blot analysis revealed that NHE6 protein was newly expressed in the corrected line (Table 1 and **Supplementary Fig. 13b**). Two subclones from a second corrected line were also generated for this mutation, and NHE6 protein was newly expressed in these corrected subclonal lines as well (**Supplementary Fig. 14**).

Next, we generated induced mutations in a founder iPSC line derived from fibroblasts of a typically developing adult male and in the HEK293T cell line, which was derived from female fetal kidney epithelia^28^. For the iPSC lines, clones were generated in which different indel mutations were present in exon 2 of *SLC9A6*; for the HEK293T lines, clones were generated in which different indel mutations were present in exon 1 of *SLC9A6* (Table 1**; Supplementary Fig. 13c;** and **Supplementary Fig.15a,b**). NHE6 protein was not detected in these mutant lines, as shown by immunoprecipitation of NHE6 followed by western blot analyses (**Supplementary Fig. 13d** and **Supplementary Fig.15c**). Thereby, we have now established multiple lines of cells with distinct NHE6 mutations, with robust controls. These cell lines include: patient-derived iPSCs with NHE6 mutations, unaffected sibling control iPSCs, isogenic corrected iPSCs, founder iPSCs and isogenic targeted NHE6-mutant iPSCs, and HEK293T control and targeted NHE6-mutant cells (Table 1; and **Supplementary Fig. 15a-c**).

### NHE6 mutation leads to endosomal over-acidification that is reversed by genome correction in patient lines

Loss-of-function mutations in NHE6 and homologs have been associated with a unique endosomal phenotype. As NHE6 is presumed to be an abundant exchanger that permits leak of protons from early, recycling, and late endosomes, loss of NHE6 has been shown to lead to over-acidification of endosomes^15, 29-31^. We conducted live-cell fluorescent ratiometric measurements of endosomal pH in CS and control iPSCs using fluorescein isothiocyanate (FITC, pH sensitive) and Alexa-546 (pH insensitive) conjugates of transferrin together with an endocytosis and flow cytometry-based analysis protocol^32^ (Fig. 3a,b**; Supplementary Table 2;** and **Supplementary Fig. 16**). We initially measured intra-endosomal pH in iPSCs derived from a CS patient as compared to iPSCs derived from the genetically related non-carrier brother. A subset of patient lines showed a strong reduction of intra-endosomal pH, as discussed below. However, not all patient-derived lines demonstrated strong reductions in intra-endosomal pH relative to their respective paired lines derived from a biologically related control (**Supplementary Table 2** and **Supplementary Fig. 16**). To assess for the possibility that variation in the strength of the effect on intra-endosomal pH across patient lines was due to background genomic variation, we quantified intra-endosomal pH in our isogenic iPSC resource wherein multiple distinct NHE6 mutations were induced in a single founder line. Here we again noted that some mutant clones demonstrated a greater reduction in intra-endosomal pH than other mutant lines relative to the isogenic control (**Supplementary Table 2** and **Supplementary Fig. 16**). This finding was again reproduced in control and induced-mutation HEK293T cell lines wherein one induced mutations showed a strong effect and the second lines showed a more modest effect (**Supplementary Table 2** and **Supplementary Fig. 15d,e**). To further address the variation in this phenotype seen in the iPSC lines, we conducted a large number of biological replicates across all lines with multiple subclones per line studied. Using mixed effect regression modeling across all the data to account for important variables (such as mutation, subclones, number of passages), we see statistically significant effects of NHE6 mutation driving reductions in intra-endosomal pH. For the family analysis with biologically-related brothers, we see an average control intra-endosomal pH of 6.27 with a decrease in patients of approximately 0.08 pH points (*P* = 0.004, 95% confidence interval reflecting decreases from 0.027 to 0.142 pH points). Additionally, across all of the isogenic lines with induced mutations (iPSCs and HEK293T), we observed a statistically significant decrease of 0.09 pH points (*P* = 0.01, 95% confidence interval reflecting decreases from 0.02 to 0.15 pH points).

**Figure 3.**
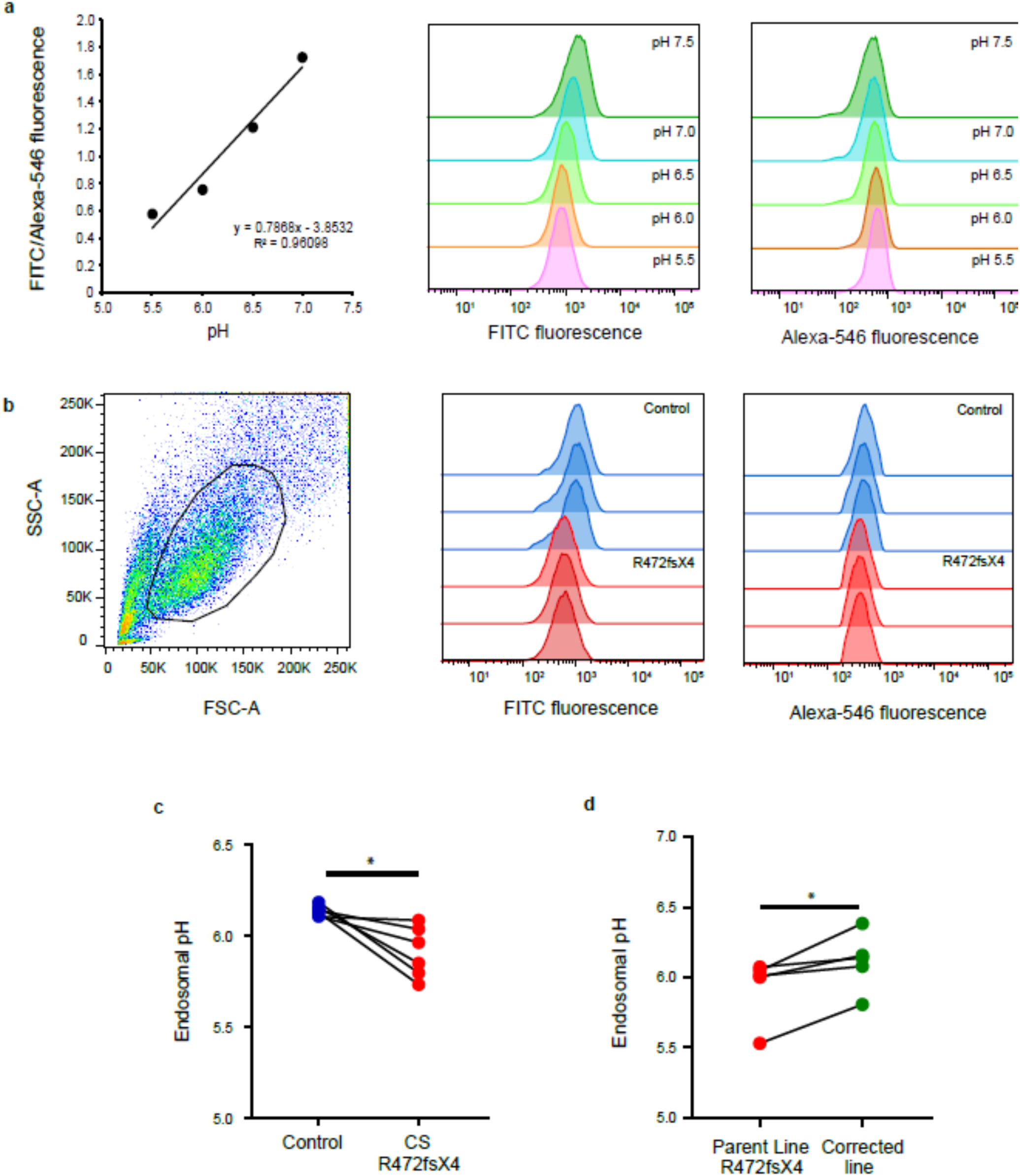
Endosomal pH defects in CS patient-derived lines can be restored in genome-corrected iPSCs. (**a**) Endosomal pH was measured using flow cytometry of iPSCs incubated with pH-sensitive (FITC) and pH-insensitive (Alexa-546) conjugates of transferrin. pH was calculated by measuring the ratio of mean FITC to mean Alexa-546 fluorescence after generating a pH standard curve. The left panel shows the calibration standard curve and equation used to calculate the endosomal pH; middle and right panels show the flow cytometer profiles for the fluorophores under each buffer condition. (**b**) Sample flow data for c.1414dupA/R472fsX4 and control iPSC lines. The left panel shows the gated population analyzed; middle and right panels show three experimental replicates for CS and control iPSC lines for the FITC (middle) and Alexa-546 (right) channels. (**c**) Graph depicting endosomal pH, as measured in c.1414dupA/p.R472fsX4 and paired control iPSCs. *n* = 6, paired t-test, *P* = 0.017. (**d**) Graph depicting endosomal pH, as measured in c.1414dupA/p.R472fsX4 parent iPSCs and isogenic, genome-corrected iPSCs. *n* = 5, paired t-test, *P* = 0.0308.

We assessed variability in the effects of NHE6 mutations on intra-endosomal pH even further by analyzing intra-endosomal pH in a patient-derived c.1414dupA/p.R472fsX4-mutant subclonal iPSC line as compared to a subclonal line of isogenic corrected iPSCs. Here the patient-derived line with the mutation was more acidic by 0.23 pH points on average as compared to the unaffected brother’s iPSCs (*P* < 0.02) (Fig. 3c). Importantly, in the genome-corrected iPSCs, endosomal pH was increased by 0.18 pH points on average as compared to the c.1414dupA/p.R472fsX4 parent line (*P* = 0.0308) (Fig. 3d). Here too, to augment rigor, we studied intra-endosomal pH across all available rescued subclonal lines and utilizing a mixed effect regression analysis, we found a statistically strong result reflecting an increase in intra-endosomal pH with 95% confidence intervals of an increase from 0.07 to 0.14 pH points across all subclonal lines studied (*P* < 0.001). Therefore, this rescue experiment attributes this decrease in intra-endosomal pH in the patient directly to the absence of NHE6 protein. Furthermore, statistical analyses across the large number of mutations and subclonal lines studied supports a statistically significant effect of NHE6 mutations on intra-endosomal pH.

### Transcriptome analysis shows down-regulation of V-ATPase genes with loss of NHE6

To gain insight into gene expression changes that might result from and/or compensate for NHE6 mutations, we used NanoString technology to assess differential gene expression between CS and control cells, in both iPSC cultures and neuronal cultures (**Supplementary Tables 3-21 and Supplementary Figs. 17>-25**). We developed a customized NanoString gene panel (384 genes) for RNA quantification, that included 74 genes relating to neuronal development, and 278 genes that we hypothesized may be affected by loss of NHE6. The customized panel includes genes relating to: endosome biology, including endosome trafficking, endosome sorting, chloride channels, and the V-ATPase subunits; autophagy and lysosome biology, including nutrient sensing, lysosome biogenesis, and lysosomal stress; neurodegeneration, including amyloid precursor protein processing and tau; signaling through the tyrosine kinase and IGF pathways; and the NHE family members (**Supplementary Table 3;** see Methods for the rationale regarding selection of genes to include in the customized panel). Initially, iPSCs from three patient-derived mutations and their matched related controls were compared. Analysis of differential gene expression between CS and control iPSCs in 1) a grouped analysis of all CS lines and all controls (**Supplementary Tables 14 and 15**) and 2) in patients and controls within individual families demonstrated several significant differences (**Supplementary Fig. 23** and **Supplementary Tables 16-1**). Interestingly, outside of *SLC9A6*, none of the other NHE family members showed significant changes in gene expression (**Supplementary Fig. 24**). In the c.1414dupA/p.R472fsX4 iPSC lines (the line with the greatest effect on intra-endosomal pH), genes significantly downregulated included genes associated with lysosomal storage disorders, lysosomal function, and the V-ATPase (**Supplementary Table 17**). In particular, 6 out of the 13 subunits of the V-ATPase were significantly downregulated in c.1414dupA/p.R472fsX4 iPSCs, the line that shows the strongest effect with regard to intra-endosomal pH (**Supplementary Fig. 25**). This is supportive of a compensatory mechanism that might be in place, particularly in c.1414dupA/p.R472fsX4 iPSCs, to maintain pH homeostasis, given the evidence of reduced endosomal pH (**Fig. 3c,d**).

We further analyzed the iPSC NanoString data (318 genes in 12 control and 11 CS iPSC samples) using Weighted Gene Co-expression Network Analysis (WGCNA)^33^ to identify a network of highly correlated genes in CS and control iPSCs (Fig. 4 and **Supplementary Table 22**). The relationship of each gene with disease status (iPSC/CS trait) was calculated by Pearson’s correlation. We used the Hybrid Dynamic Tree Cut method and signed networks to generate a network dendrogram that contained three color-coded gene-connected modules^34,35^ (Fig. 4a-c). The power value was set at 12 in this analysis. The largest module (shown in turquoise) contains 114 genes, only 2 out of 19 of which are *VATPaseV0* and *VATPaseV1* genes. Importantly, the second largest module (shown in blue) contains 75 genes, including both *SLC9A6* and interestingly, 11 out of 19 *VATPaseV0* and *VATPaseV1* genes. Gene significance (GS) represents the biological significance level of a gene to the disease status. Higher GS (up to 1) indicates higher biological significance. The blue module shows the highest GS as compared to the turquoise and gray modules (Fig. 4d). Genes with high connectivity within a module are called “hub genes”, indicating tightly connected and co-expressed genes in the module. Notably, the hub genes in the blue module show higher GS compared to the non-hub genes (*P* = 7.8 x 10^-5^), whereas the turquoise module shows lower significance (*P* = 0.0005), and the gray module does not show any significance (*P* = 0.72) (Fig. 4e). Notably, the enrichment of the V-ATPase genes in the blue module with SLC9A6 was highly significant above what would be predicted by chance alone (*P* = 0.0006, Hypergeometric test, Reject “coincidence” hypothesis) (**Supplementary Table 23**). The correlation between *SLC9A6*, *VATPaseV0* genes, and *VATPaseV1* genes was significant within the selected module. A VisANT-like circle plot was generated based on the intra-modular connectivity of all genes in the blue module. The range of genes represents the gene connectivity order in its related module as well as node size, and lines represent connections between two nodes (Fig. 4f). This analysis indicates the significant difference between control and CS iPSCs in gene connectivity and hub gene status in the blue module, which includes both NHE6 and V-ATPase genes. These data further underscore the role of NHE6 in the biological process involving regulation of intra-organellar pH.

**Figure 4.**
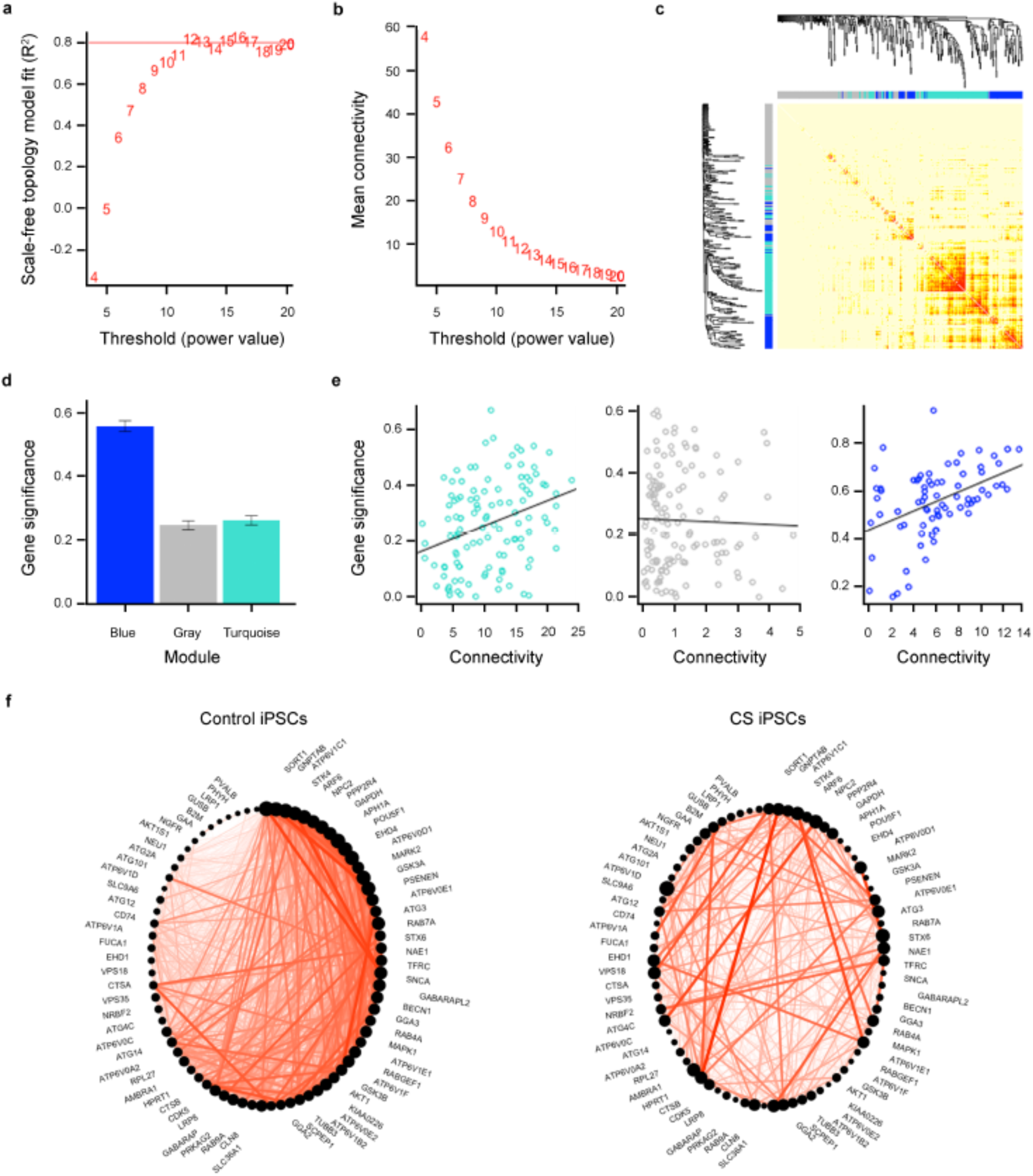
WGCNA-based network construction and module detection in CS iPSCs. (**a**) As the power value increases the scale in network reaches a plateau, indicating independence of the network. The scale in network reaches a plateau starting at 12. (**b**) As the power value increases the number of highly connected genes decreases. Higher power values separate highly connected genes from other less connected genes. (**c**) Heatmap and dendrograms based on hierarchical clustering on the dissimilarity of topological overlap measurement. Two modules, blue and turquoise, are detected. Genes not assigned to a detected module are combined together and assigned to the gray module. (**d**) Graph depicting the gene significance of the indicated modules. Gene significance is defined as the minus log of the *P*-value for Pearson’s correlation of a gene to iPSC/CS disease status. Module gene significance is defined as the average absolute gene significance of all genes in that given module. The blue module has a high positive average gene significance (*P* = 2.1 x 10^-26^, Kruskal-Wallis Test). (**e**) Graphs depicting intra-modular connectivity vs. gene significance based on performance of linear regression modeling. The Correlation and *P*-value for each module are as follows: Turquoise module (Correlation = 0.32, *P* = 0.0005), gray module (Correlation = −0.032, *P* = 0.72), and blue module (Correlation = 0.44, *P* = 7.8 x 10^-5^). (**f**) Circle plots of the connectivity of genes in the blue module for control iPSCs (left) and CS iPSCs (right). The size of a dot is indicative of a gene’s connectivity within the module; the bigger the dot, the higher the connectivity. Red lines indicate the connection between two genes. Note that the dot for *SLC9A6* is smaller in the circle plot for control iPSCs as compared to the dot for *SLC9A6* in the circle plot for CS iPSCs, indicating greater connectivity with other genes in CS iPSCs.

### Postnatal microcephaly is associated with defects in neuronal arborization in patient-derived neurons and is not attributable to differences in proliferation or cell death

Postnatal microcephaly, such as observed in CS patients, may result from deficiencies of neuronal arborization in the neocortex, as opposed to differences in progenitor cell proliferation or cell death; in contrast, primary microcephaly generally occurs as a result of defects in cortical progenitor neurogenesis, cell fate determination, or pathways leading to cell death^4^. To determine the potential mechanisms of postnatal microcephaly in CS, we differentiated CS and control iPSCs to an excitatory cortical neuronal fate using a monolayer differentiation protocol involving dual SMAD inhibition^36^ (**Supplementary Fig. 17**). To assess the ability of our CS and control iPSCs to differentiate to a cortical fate, we analyzed the expression of genes relating to neuronal development and differentiation – before and after differentiation – using NanoString-based analyses together with a customized panel of 74 marker genes (**Supplementary Table 3**). The selected genes included markers for neuronal progenitors, neurons, cortical layers, synapses, interneurons, glia, midbrain, mesoderm, endoderm, and pluripotency. Hierarchical clustering revealed that changes in gene expression discriminated neuronal cultures from iPSC cultures (**Supplementary Fig. 18**). There were numerous differentially expressed genes between the iPSC state and neuronal state (combining mutant and control lines together) (false discovery rate (FDR)-adjusted *P*-value < 0.05) (**Supplementary Fig. 19** and **Supplementary Tables 4 and 5**), reflecting the strength of our differentiation methods.

In order to examine the possibility that differences in neuronal progenitor proliferation, cell fate determination, or cell death could contribute to CS-associated microcephaly, CS and control neuronal cultures were examined using immunocytochemistry and RNA expression profiling. By immunocytochemistry, CS and control neuronal cultures had comparable percentages of cells expressing markers for neuronal progenitors (PAX6), neurons (MAP2), and deep-layer cortical projection neurons (TBR1, CTIP2) (**Supplementary Fig. 20**). We further examined the cell fate of CS and control neuronal cultures using our NanoString-based panel of mRNAs related to neuronal cell fate determination. In neuronal cultures, cells of both genotypes analyzed in batch format (**Supplementary Tables 6 and 7**) or within families (**Supplementary Tables 8-13**) expressed comparable and high levels of neuronal progenitor, neuronal, and cortical markers, and very low levels of mesodermal, endodermal, glial, and midbrain markers (**Supplementary Fig. 21**). Therefore, these analyses demonstrate that cortical cell fate determination in these cultures does not appear to be affected by NHE6 mutations in patient-derived cells. We also ruled out differences in neuronal cell proliferation or cell death between CS and control iPSC-derived neurons. The number of mitotically cycling cells positive for Ki67 was not statistically different between CS and control cultures at day 35 after neuronal induction. Similarly, there was no difference in the number of apoptotic cells between CS and control neuronal cultures (**Supplementary Fig. 22**). We conclude that mechanisms frequently involved in primary microcephaly such as alterations in cell fate determination, progenitor proliferation, or cell death are unlikely to contribute to CS-associated secondary or postnatal microcephaly.

In prior studies in NHE6-null mice, we identified abnormalities in neuronal arborization both *in vitro* and *in vivo*. Reduced neuronal arborization may contribute to postnatal microcephaly in CS, particularly in light of our data above showing no differences in cell proliferation or death. Therefore, to expand on our prior findings in mouse but now in neurons from human patients, we examined neurite outgrowth and arborization in neurons differentiated from CS patient iPSCs. Importantly, across all CS mutations examined, using a variety of different approaches, we saw prominent defects reflecting abnormal neurite growth and arborization in CS. For example, we conducted time-lapse live imaging analysis of CS neurons plated on laminin stripes (Fig. 5a). The neurons aligned along the laminin stripes and extended and retracted their neurites. We found significant decreases in the elongation rates and retraction rates, in comparison to their respective controls, for specific CS lines (Fig. 5b). Additionally, using Sholl analysis to assess arborization, we observed strong defects in CS neurons regardless of the CS mutation studied (**Supplementary Fig. 26**). We then measured the intra-endosomal pH within neurons using a fluorescence-based ratiometric assay^15,32^ and tested the hypothesis that the effect on arborization would be correlated with the quantitated decrease in intra-endosomal pH in neurons (**Supplementary Fig. 27**). This analysis determined that all mutations demonstrated prominent decreases in neuronal arborization, regardless of the nature of the mutation examined or the strength of the measured change in intra-endosomal pH. Taken together, these data from live imaging and Sholl analysis suggest that defects in arborization might underlie CS patient phenotypes such as postnatal microcephaly.

**Figure 5.**
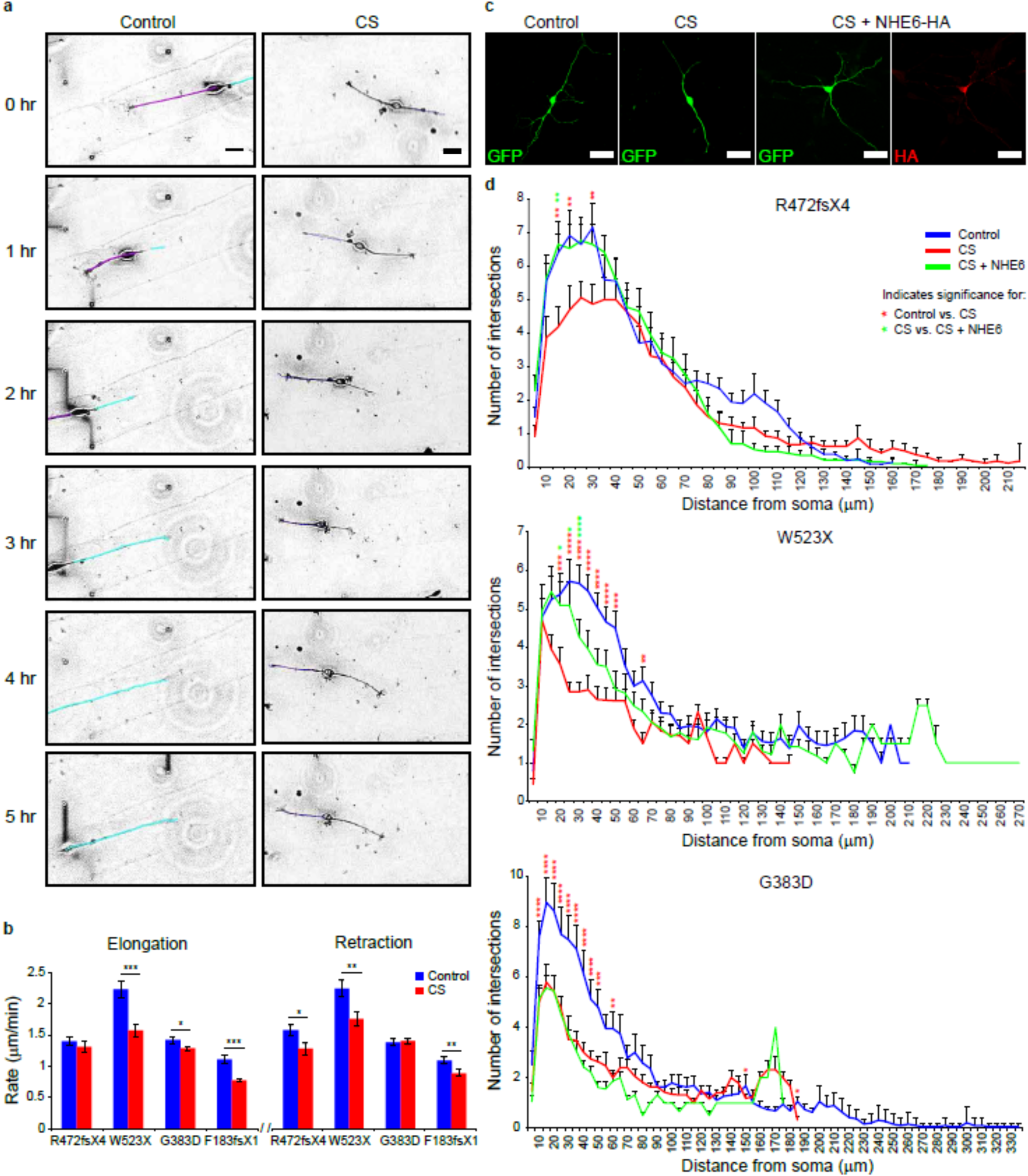
Deficits in neuronal arborization can be rescued by re-expression of NHE6 in CS lines with frameshift or nonsense mutations but not in the CS missense mutation line. (**a**) Neurite outgrowth dynamics in iPSC-derived neurons. Time-lapse images of iPSC-derived neurons plated on laminin-striped coverslips. Time-lapse images were acquired using Volocity software at four XY positions every 5 min for 12 hr. Representative cells and tracings are shown from the c.1148G>A/p.G383D mutant line and paired control line. Scale bars, 20 μm. (**b**) The rate of neurite elongation and retraction were measured using ImageJ software for the first 2 hr. Mean ± s.e.m., Welch’s t-test: * *P* < 0.045, ** *P* < 0.008, *** *P* < 0.00008. Family 1, c.1414dupA/p.R472fsX4: *n* = 14 cells/ 34 neurites/ 875 measurements. Control Family 1: *n* = 14 cells/ 33 neurites/ 841 measurements. Family 2, c.1568G>A/p.W523X: *n* = 6 cells/ 9 neurites/ 724 measurements. Control Family 2: *n* = 12 cells/ 15 neurites/ 635 measurements. Family 3, c.1148G>A/p.G383D: *n* = 53 cells/ 126 neurites/ 2,163 measurements. Control Family 3: *n* = 28 cells/ 55 neurites/ 827 measurements. Family 4, c.540_547dupAGAAGTAT/p.F183fsX1: *n* = 14 cells/ 25 neurites/ 708 measurements. Control Family 4: *n* = 8 cells/ 17 neurites/ 622 measurements. (**c**) Cell-autonomous gene replacement experiment. iPSC-derived neurons were transfected with a construct encoding for GFP alone or with two constructs together encoding for GFP + full-length human NHE6-HA and analyzed after 5 days. Representative images of each condition are shown. Scale bars, 20 μm. (**d**) Sholl analysis of control iPSC-derived neurons (blue lines), CS iPSC-derived neurons (red lines), and CS iPSC-derived neurons transfected with a construct encoding for NHE6-HA (green lines). The average number of neurite intersections with Sholl radii was plotted as a function of distance from the soma for each condition. *n* = 20-25 cells per condition. Mean ± s.e.m., two-way ANOVA with Bonferroni multiple comparisons test, * *P* < 0.05, ** *P* < 0.01, *** *P* < 0.0001, **** *P* < 0.00001. Red asterisks denote significance when comparing CS to control cells. Green asterisks denote significance when comparing CS to CS cells transfected with a construct encoding for NHE6.

### Cell-autonomous cDNA replacement rescues neuronal arborization in patient neurons with nonsense mutations but not with missense mutation

Using Sholl analysis to assess arborization, we observed strong effects regardless of the CS mutation studied. However, with regard to cell-autonomous rescue of these defects using gene replacement by transfection in patient-derived neurons, cells with the protein-truncating mutations c.1414dupA/p.R472fsX4 and c.1568G>A/p.W523X behaved alike to each other, but differently from the cells with the missense/splice mutation c.1148G>A/p.G383D. For these studies, we transfected CS and control neurons with a single construct encoding for GFP or with a construct encoding for GFP together with a construct encoding for full-length NHE6 tagged with HA (Fig. 5c). Using Sholl analysis, we found that, in all three CS mutations, there was a strong decrease in the number of times neurites intersected Sholl radii, as compared to the paired controls (Fig. 5d). Furthermore, the neurite length within each radius was decreased in all CS mutations at distances close to the soma (**Supplementary Fig. 26**). Cell-autonomous re-expression of NHE6-HA led to prominent rescue of arbor complexity (by Sholl analysis) in neurons with protein-truncating mutations; however, rescue completely failed in neurons from the complex missense/splice mutation (Fig. 5d and **Supplementary Fig. 26**). Notably, re-expression of NHE6 in the c.1148G>A/p.G383D CS line decreased the number of total branches, branchpoints, and neurite length, while these measures were increased in the rescue experiments in neurons from the patients with nonsense mutations (**Supplementary Fig. 28**).

This rescue experiment reveals important distinctions about the mechanisms of action of patient mutations and has implications for therapeutics. First, restoration of arbor complexity in c.1414dupA/p.R472fsX4 and c.1568G>A/p.W523X CS cells reveals a cell-autonomous mechanism of rescue, which is strongly consistent with our prior data indicating that these mutations are complete nulls. Second, the failure of NHE6 re-expression to rescue arbor complexity in c.1148G>A/p.G383D CS cells suggests that this NHE6 missense mutation may behave as a dominant-negative by forming heterodimers with newly expressed NHE6. Thereby, while gene replacement by re-expression might be a plausible therapeutic strategy in complete null mutations, it is unlikely to be a successful therapeutic strategy in the context of complex missense mutations or other similar types of mutations.

### G383D missense mutant protein has both loss-of-function and dominant-negative properties

The failure to rescue arborization with gene replacement (cDNA transfer) in G383D-mutant neurons suggests that the remnant endogenously expressed protein may exert a dominant-negative function through formation of non-functional dimers. We addressed this question mechanistically, by testing if exogenously expressed G383D protein could form dimers with exogenously expressed wild-type NHE6 protein in HEK293T cells. Similarly to endogenously expressed G383D in iPSCs (Fig. 1c), we find that exogenously expressed G383D has reduced stability as compared to wild-type NHE6 protein (Fig. 6a,b and **Supplementary Fig. 29**). We also find that exogenously expressed G383D protein can form homodimers with itself, and heterodimers with exogenously expressed wild-type NHE6 protein (Fig. 6a,b). However, dimer formation containing G383D protein is decreased relative to controls.

**Figure 6.**
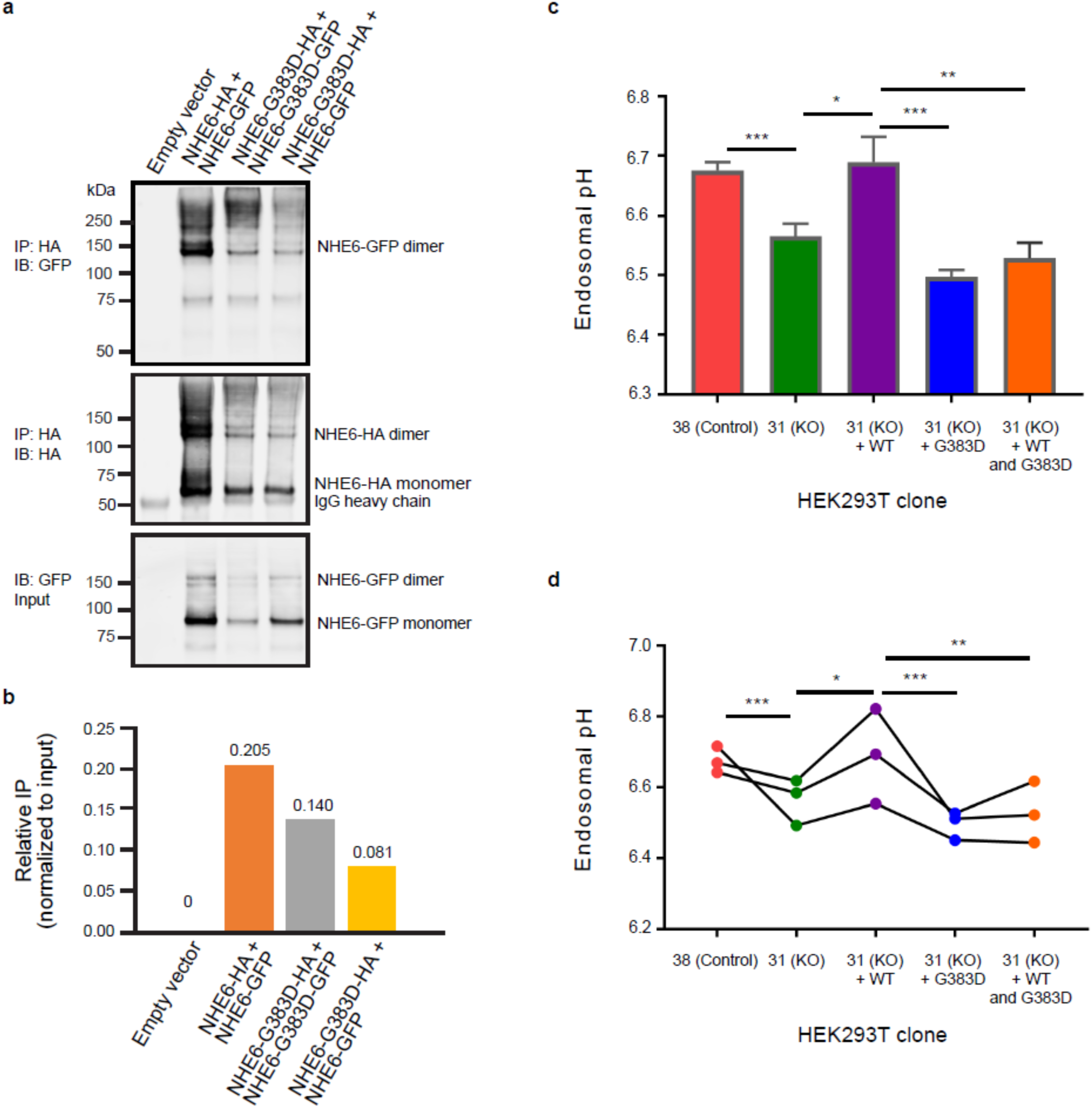
The G383D mutation in NHE6 decreases the formation of NHE6 dimers and prevents full rescue of defects in endosomal pH. (**a**) Expression of the NHE6-G383D mutant decreases the ability of NHE6 to form dimers. Lysates from HEK293T cells expressing the indicated tagged forms of NHE6 were immunoprecipitated using an anti-HA antibody and subsequently analyzed by western blot using antibodies against GFP (top panel) and HA (middle panel). Protein input was determined by western blot of lysates that had not undergone immunoprecipitation for HA-tagged protein using an antibody against GFP (bottom panel). (**b**) Graph depicting the relative amount of immunoprecipitated GFP-tagged protein based on results shown in panel (**a**). Immunoprecipitates were normalized to protein expression levels. IP = immunoprecipitation. (**c**) Graph depicting endosomal pH, as measured in HEK293T control cells (clone 38, Control) and HEK293T hSLC9A6 knockout cells (clone 31, KO). The hSLC9A6 knockout cells were analyzed as the clonal line or in the context of expression of NHE6-WT-mCherry (+ WT), NHE6-G383D-mCherry (+ G383D), or NHE6-WT-mCherry and NHE6-G383D-mCherry (+ WT and G383D). Mean ± s.e.m., *n* = 3, unpaired t-test, *** *P* = 0.00043 (clone 38 vs. clone 31), * *P* = 0.017 (clone 31 vs. clone 31 + WT), *** *P* = 0.00042 (clone 31 + WT vs. clone 31 + G383D), ** *P* = 0.0048 (clone 31 + WT vs. clone 31 + WT and G383D). (**d**) Plot depicting endosomal pH, based on data represented and described in panel (**c**). Each dot indicates the average of three replicas from one independent experiment.

Next, we hypothesized that G383D could act as a dominant negative and asked whether G383D could: 1) function to alkalinize endosomes and/or 2) inhibit wild type NHE6 protein function. To test these mechanisms, we generated a CRISPR/Cas9 genome edited HEK293T cell line, wherein NHE6 was mutated and in which we observed significant decrease in endosomal pH relative to control (**Supplementary Fig. 15**). Expression of a wild type NHE6-mCherry construct in HEK293T NHE6 mutant cells resulted in an alkalization of endosomes as compared to untransfected HEK293T NHE6 mutant cells (Fig. 6c,d). However, expression of an NHE6-G383D-mCherry construct in HEK293T NHE6 mutant cells did not alkalinize the endosomes (Fig. 6c,d). Importantly, co-expression of NHE6-G383D-mCherry and wild type NHE6-mCherry constructs in HEK293T NHE6 mutant cells inhibited the ability of exogenously expressed wild type NHE6 to alkalinize endosomes (Fig. 6c,d). We posit that wild type NHE6 and G383D-mutant NHE6 heterodimers are unable to conduct cation exchange, consistent with the failure of exogenously expressed wild type NHE6 to rescue neuronal arborization in G383D iPSC-derived neurons (Fig. 5d). In summary, we find that the NHE6 G383D mutant does not function in the alkalinization of endosomes as NHE6 WT does. This was predicted by the structural analysis of the exchanger domain (Fig. 2), which suggested G383D is a loss-of-function mutation. Our dimerization and pH studies also demonstrate that G383D mutant protein might function as a dominant-negative by inhibiting the wild-type NHE6 protein function upon formation of (non-functional) heterodimers. Overall, our data support a model wherein rescue strategies involving gene replacement will likely be allele specific. While they may function in cells with a simple loss-of-function mutation, cells with complex mutations such as G383D mutation do not appear to respond to this rescue strategy.

### Cell-non-autonomous rescue of neuronal arborization by exogenous trophic factor treatment (BDNF or IGF-1) in patient neurons regardless of mutation

Herein, we have established a substantial cellular resource that may be used in cell-based assays for pre-clinical development of therapeutics. We next tested if neuronal arborization could be restored in patient-derived neurons pharmacologically through treatment with exogenous BDNF or IGF-1. We tested these two growth factors as proof-of-principle molecules, as we have previously shown that BDNF signaling is diminished in the NHE6-null mouse and, furthermore, that arborization may be rescued by exogenous application of excess BDNF in mutant mouse primary neuronal cultures^15^. Further still, IGF-1 is also a trophic factor that augments neurite growth and is currently in clinical trials for a number of neurodevelopmental disorders^37,38^. When CS and control neurons were treated for three days with exogenous addition of either BDNF or IGF-1, we observed substantial rescue of the arborization defect in all family lines tested, regardless of mutation (Fig. 7a,b). Treatment with BDNF or IGF-1 increased the mean length per neurite as well as the number of branchpoints per neurite in all CS lines, as compared to untreated CS neurons (Fig. 7b). Therefore, neurite morphology defects in CS cultures may be restored by exogenous trophic factor treatment. Together, these results suggest that independent exposure to BDNF or IGF-1 can rescue the neuronal arborization phenotype in CS neurons and warrant further exploration in existing animal models and other pre-clinical studies. Unlike the gene rescue experiments above, these exogenous trophic factors demonstrate efficacy *in vitro* across patient-derived neurons with diverse mutations, regardless of distinct underlying biochemical mechanisms.

**Figure 7.**
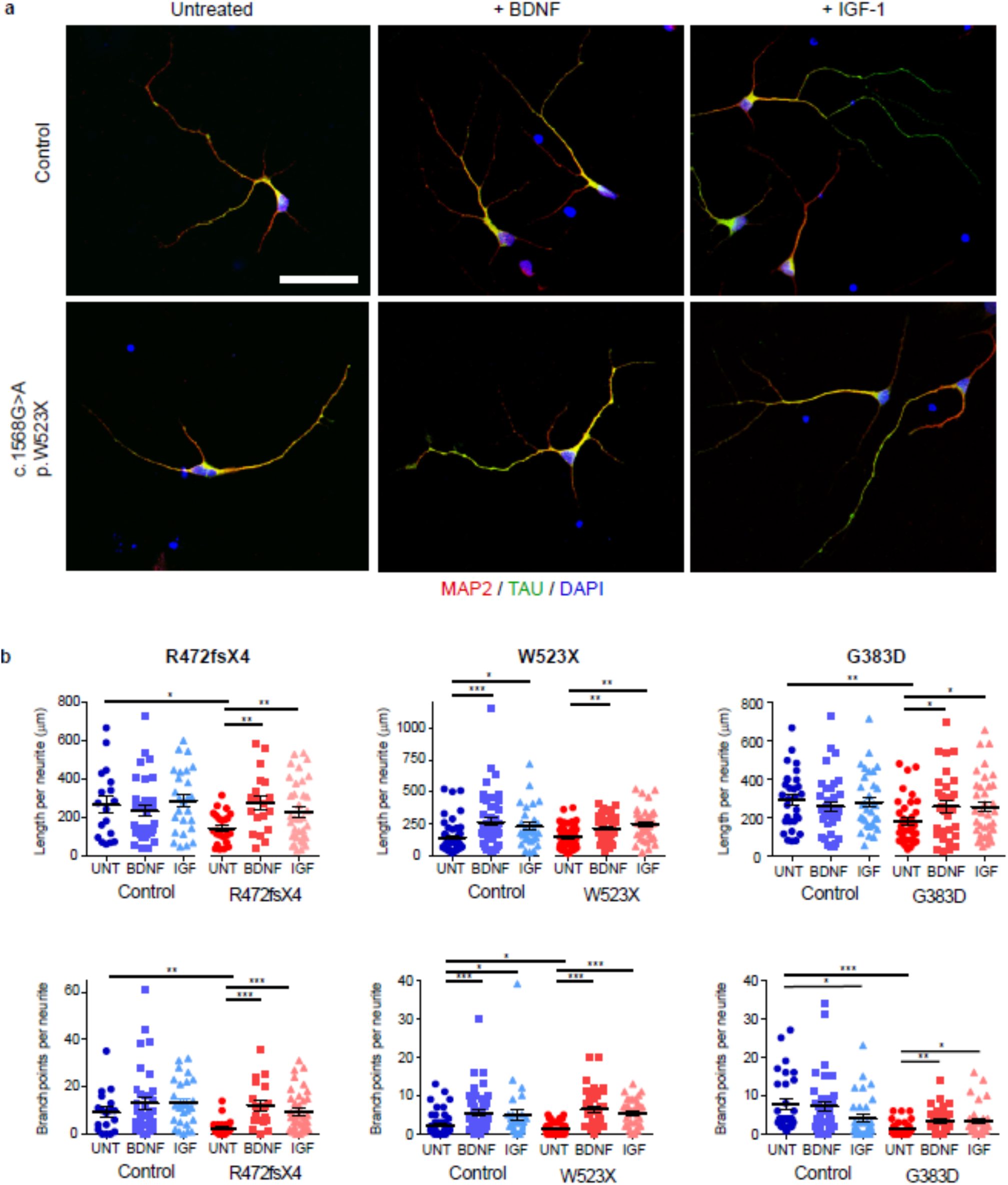
BDNF and IGF-1 treatments rescue deficiencies in neuronal arborization regardless of mutation type. (**a**) Defects in arborization are seen in all CS lines, as compared to paired controls. Immunostaining for MAP2 and TAU in iPSC-derived neurons treated with BDNF or IGF-1. Representative images are shown from the c.1568G>A/p.W523X CS line and the paired control line. Scale bar, 50 μm. (**b**) Average length and number of branchpoints per neurite were measured using Neurolucida tracing software. Family 1 c.1414dupA/p.R472fsX4: *n* = 24 (UNT),19 (BDNF), 36 (IGF) neurites. Control Family 1: *n* = 18 (UNT), 35 (BDNF), 26 (IGF) neurites. Family 2 c.1568G>A/p.W523X: *n* = 53 (UNT), 37 (BDNF), 37 (IGF) neurites. Control Family 2: *n* = 54 (UNT), 48 (BDNF), 28 (IGF) neurites. Family 3 c.1148G>A/p.G383D: *n* = 32 (UNT), 31 (BDNF), 35 (IGF) neurites. Control Family 3: *n* = 30 (UNT), 35 (BDNF), 32 (IGF) neurites. Mean ± s.e.m., Welch’s t-test, * *P* < 0.05, ** *P* < 0.005, *** *P* < 0.0005. Neurites stained with either TAU or MAP2 were analyzed; no distinction was made between dendrites or axons.

## Discussion

Despite the increasing reports of CS diagnosis in the clinic, much remains to be understood concerning the nature of the cause and treatment of CS. The generation of cortical-like human forebrain neurons from CS patient-derived iPSCs presents a new opportunity to study human neurodevelopment in CS. In this study, properties of neurons that contribute to deficits in circuitry development and several routes of intervention to restore these deficits to levels similar to those in control neurons without mutations were examined. We show that human neurons derived from CS patients have reduced complexity of neuronal arbors regardless of the nature of the mutations studied here. These deficits in neuronal growth and arborization may underlie the postnatal microcephaly reported in these CS patients and others^2^. We demonstrated that the neuronal arborization defect could be rescued by re-expression (i.e., exogenous expression) of NHE6 in cells with loss of protein, but not in cells from a patient with a complex missense/splice mutation, the G383D mutation that harbors dominant-negative activity. By contrast, the neuronal arborization defect could be rescued *in vitro* by exogenous addition of BDNF or IGF-1 regardless of mutation. BDNF agonists have been subject to efforts in pre-clinical studies of neurological disease with mixed results^39^. Alternatively, IGF-1 has been shown recently to be a potential therapy for the autism-associated syndromes Rett syndrome and Phelan-McDermid syndrome. Pre-clinical studies in Rett syndrome showed that treatment of iPSC-derived neurons with IGF-1 increased levels of glutamatergic synapses^40^. Furthermore, treatment of a Rett syndrome mouse model with IGF-1 rescued brain size, synapse density, and excitatory synapse transmission to control levels^41^. In a phase I clinical trial for IGF-1 in Rett syndrome, patients showed improved symptoms of anxiety and breathing difficulty^37^. In addition, IGF-1 treatment of iPSC-derived neurons from Phelan-McDermid syndrome patients improved deficits in mature excitatory synapses, as compared to controls^42^. A preliminary clinical trial of IGF-1 in Phelan-McDermid syndrome showed improvement of social withdrawal and restricted behaviors^38^. Additionally, a phase II clinical trial for IGF-1 is in progress with respect to a potential treatment for autism spectrum disorder (NCT01970345), and phase II trials have been completed with respect to the use of an IGF-1 analog (NNZ-2566) as a potential treatment for Fragile X syndrome (NCT01894958) and Rett syndrome (NCT02715115). Careful attention assessing the efficacy of IGF-1 in these other disorders may have relevance to therapeutic development in CS. Additionally; further research into efficacy of IGF-1 treatment may also be warranted in the CS mouse models in vivo.

Our data also show allele-specific responses to treatments with regard to gene replacement (i.e., cDNA transfer). One of the strengths of working with patient-derived iPSCs is that this methodology allows for a full determination of the effects of genetic mutation on protein coding and function based in gene expression from the endogenous locus. Studies to address these sorts of questions are often done via overexpression, which is subject to artifact, and NMD of mRNA is often overlooked. In CS, the majority of mutations introduce nonsense mutations^2^. Here we demonstrate that these mutations lead to NMD and complete or near complete loss of mRNA, and thereby, protein, i.e., a truncated protein is unlikely to be produced in large quantities. Additionally, we determine the defects resulting from an apparent missense mutation (G383D) to be quite complex, involving a combination of splicing and NMD mechanisms, as well as generation of a low level of non-functional protein that appears to act in a dominant-negative fashion. This mutation specifically fails to be rescued by way of gene replacement, yet it is rescued using cell-non-autonomous strategies (i.e., BDNF and IGF-1 addition). Allele-specific mechanisms of treatment response may well be more the rule than the exception in genetic disease, as has been studied in-depth in cystic fibrosis^19,20^.

Results of our study of intra-endosomal pH across a diverse range of patient-derived and induced (through genome editing) mutations also provide new insights into the function of NHE6 as an endosomal proton/cation exchanger. Prior data from our laboratory and others have demonstrated reductions in organellar luminal pH in the context of NHE6 or Nhx1 mutation using inbred, laboratory strains such as mouse or yeast, respectively^15,43^. However, here, using patient-derived lines from human populations with a high degree of background genomic variation, we observe a range in the strength of the effect on intra-endosomal pH. There may be several explanations for this, among them are that cells in culture undergo genetic changes that compensate for the effect of the NHE6 mutation. While we do see a weak trend of cell passages on reversion of intra-endosomal pH towards control (*P* = 0.08 in the mixed effect model; **Supplementary Table 2**), there are other possible interpretations. For example, while it is likely that the NHE6 mutations affect its ability to function as a pathway for proton leak, the defective mechanisms may also involve actions that are not best exemplified by measurable changes in intra-endosomal pH. Our transcriptome data do support a link between loss of NHE6 and changes in expression of V-ATPase genes, thereby further implicating a role for NHE6 in cross-membrane proton transport. At the same time, the defects observed in neuronal development (i.e., arborization) do not appear to clearly correlate with the strength of the effects on acidification of the endosomal lumen. These findings suggest a potential compensation of low intra-endosomal pH in cultured cells. Overall, the weight of the data here suggests that NHE6 also functions in a process yet to be determined (not apparently attributable solely to alterations in intra-endosomal pH) that perturbs neuronal development. Also, these results raise the question as to whether alkalinizing agents (such as V-ATPase inhibitors or weak bases) will be effective in treating CS, as over-acidification of the endosomal lumen may not be a universal feature. Our results also highlight the relevance of the contributions of specific mutations and genomic background to the disease, even in a monogenic disorder.

Finally, a major strength of our study is the establishment and validation of these iPSC resources for the study of cellular mechanisms of disease and pre-clinical drug development. Our resource, which will be widely shared, has established patient-derived cells with both related (unaffected brother) and isogenic (from mutation correction) controls, as well as induced mutations in a normal male founder iPSC line and in HEK293T cells. Through the results of our studies described here, we present merits and potentially some limitations (i.e., background genetic variation) in using distinct iPSC systems; however, we would also like to highlight the clear value of having access to multiple mutations, controls, and subclonal lines for use in carrying out studies. We are hopeful that the study results presented here and the extensive human cell resources developed will be of benefit to the research community and catalyze the development of therapeutic treatments for CS.

## Methods

### Generation of iPSCs and maintenance

The Institutional Review Boards at Brown University and Lifespan Healthcare approved all human subjects research protocols. Peripheral blood donated with informed consent from either CS patients or unaffected genetically related brothers was mixed with heparin (AAP Pharmaceuticals) to a final concentration of 23 U/mL heparin. The CS-cohort iPSC lines used in this study were generated by the Cincinnati Children’s Hospital Medical Center Pluripotent Stem Cell Facility. The protocol for generation of iPSCs from PBMCs was adapted from Kunisato, *et al.*^22^. PBMCs were enriched by layering blood diluted 1:1 with phosphate-buffered saline (PBS) + 2% fetal bovine serum (FBS) over Ficoll-Paque PLUS (StemCell Technologies). After centrifugation at 400 x *g* for 30 min, the upper plasma layer was discarded and the mononuclear cell layer was removed. PBMCs were washed in PBS + 2% FBS and plated at 5 x 10^6^ cells per well of a 6-well ultra-low attachment plate in media containing: 1X Ex-Vivo 15 media (Lonza), 10% dialyzed fetal calf serum (dFCS) (Hyclone), and 100 ng/mL human steel factor (SCF), 100 ng/mL human thrombopoietin (TPO), 100 ng/mL interleukin 3 (IL-3), 20 ng/mL interleukin 6 (IL-6), 100 ng/mL Fms-related tyrosine kinase 3 ligand (Flt3L), 100 ng/mL granulocyte-macrophage colony-stimulating factor (GM-CSF), and 50 ng/mL macrophage colony-stimulating factor (M-CSF) (PeproTech). Two days later, adherent and suspension PBMCs were collected and transduced with vesicular stomatitis virus-G (VSV-G) pseudotyped lentivirus (pRRL.PPT.SF.hOct3/4.hKlf4.hSox2.hMyc.i2dTomato.pre)^44^ expressing *SOX2, OCT4, KLF-4,* and *c-MYC* from a polycistronic mRNA at a multiplicity of infection of 25. Cells were incubated with 1,000X polybrene (Sigma) and lentivirus for 3 hr at 37°C in one well of a 24-well dish and then re-plated into one well of a 6-well ultra-low attachment dish. Two days after transduction, cells were re-plated onto mouse embryonic fibroblasts (plated 1 day prior) in media containing 1X Ex-Vivo 15 media (Lonza), 10% dFCS (Hyclone), and 100 ng/mL SCF, 100 ng/mL TPO, 100 ng/mL IL-3, 20 ng/mL IL-6, and 100 ng/mL Flt3L (PeproTech). A half change of media was performed four days after transduction followed by a full change on day 6 to hESC media containing: 1X DMEM/F12 (Sigma), 4 ng/mL human basic fibroblast growth factor (bFGF) (PeproTech), and 20% Knock-out Serum Replacement (KOSR), 50 U/mL penicillin-streptomycin, 2 mM L-glutamine, 100 μM β-mercaptoethanol, and 100 μM nonessential amino acids (Invitrogen). Cultures were fed daily with hESC media and were monitored for the development of iPSC colonies. Colonies were expanded 20-30 days after transduction by manual dissection. iPSC lines were negative for mycoplasma as tested by Lonza MycoAlert. iPSC clonal lines were cultured in feeder-free conditions on Matrigel (Corning 354277)-coated plates and maintained in mTeSR1 media (StemCell Technologies 85850). Enzyme-free passaging was performed using ReLeSR (StemCell Technologies 05872).

### Alkaline phosphatase activity

Pluripotency was assessed using the alkaline phosphatase staining kit (Millipore SCR004) as specified by the manufacturer. Briefly, putative iPSC cultures were fixed with a 3.7% formaldehyde solution in PBS for 2 min and permeabilized in TBS-T buffer containing 20 mM Tris-HCl, pH 7.4, 0.15 M NaCl, and 0.05% Tween-20. Alkaline phosphatase activity was detected following incubation in Naphthol:Fast Red Violet solution in H_2_O (mixed at a ratio of 2:1:1) for 20 min in the dark at room temperature. Substrate was replaced with TBS-T for imaging and identification of alkaline phosphatase-positive colonies.

### Karyotype

Standard G-banded metaphase spreads were prepared and interpreted by a cytogeneticist in the Cincinnati Children’s Hospital as previously described^45^.

### Fluorescence-activated cell sorting (FACS) analysis for iPSC characterization

Antibodies against TRA-1-60 and SSEA4 were used for flow cytometric analysis of pluripotency marker expression. Briefly, iPSCs were dissociated to single cells with Accutase (Thermofisher 00-4555-56) and 0.5-1.0 x 10^6^ cells were incubated on ice for 30 min with FITC mouse anti-human TRA-1-60 (BD Pharmingen 560876; 62.5 μg/reaction) or PE mouse anti-human SSEA4 (BD Pharmingen 560128; 7.5 μg/reaction) antibodies diluted in wash buffer (PBS containing 0.2% bovine serum albumin (BSA) and 0.05% sodium azide). To exclude dead cells, 7AAD was included at a concentration of 2.5 μg/mL. Unstained cells and cells incubated with the isotype control antibodies FITC mouse IgM (BD Pharmingen 553474) and PE mouse IgG3 (BD Pharmingen 559926) were used to define positive TRA-1-60 and SSEA4 populations.

### Embryoid body differentiation

For embryoid body (EB) formation, iPSCs were pre-differentiated for 3 days in EB media consisting of DMEM containing 20% FCS and 1X nonessential amino acids. Pre-differentiated cultures were then incubated with 1 mg/mL Dispase (StemCell Technologies 07913) for 5 min and washed 3 times with DMEM/F12 50:50 media. Following addition of EB media, a pulled glass pipet was used to score colonies. Colony pieces were then lifted using a cell scraper, transferred into a 6 well ultra-low attachment dish in EB media, and incubated overnight at 37°C with gentle rotation. The following day, EBs were transferred to a 15 mL Falcon tube, gently triturated to remove debris from the edges of the EBs, and re-plated in an ultra-low attachment dish in EB media. After 4 days in suspension culture, EBs were plated in EB media on 0.1% gelatin-coated cell culture plates for an additional 10 days until harvest and analysis. EB media was changed every 2 days.

### qRT-PCR for iPSC characterization

Total RNA was harvested from undifferentiated and EB-differentiated hPSCs using Trizol (Thermofisher 15596018), and DNA was eliminated using the DNase-free kit (Thermofisher AM1906). RNA was reverse transcribed to cDNA using the High-capacity cDNA Reverse Transcription Kit with RNase Inhibitor (Thermofisher 4374967) following the manufacturer’s recommendations. Gene expression levels were determined using 96-well Fast Custom TaqMan Array Plates containing Taqman assays for detection of pluripotency markers (*Pou5F1*, *NANOG, GABRB3, DNMT3B, TDGF1*, and *ZPF42*) or germ layer-specific markers (endoderm: *AFP*, *ALB*, *FABP2*; mesoderm: *TNNT2, HBZ, MEOX1*; ectoderm: *PAX6, TH, KRT1*). Taqman reactions were performed in a StepOne Plus real-time PCR device following the manufacturer’s protocols and consisted of 20 ng of each cDNA in 1X Fast Universal PCR Master Mix in a final volume of 10 μL. Thermal cycling conditions were 50°C for 2 min and 95°C for 30 s followed by 40 cycles of 95°C for 3 s and 60°C for 30 s. Relative expression of each gene was calculated for each sample using the ΔΔCt method. For pluripotency gene expression analysis (**Supplementary Fig. 8**), mean expression (± s.d. from 2 replicates) of each gene is shown relative to the expression of the same gene in H1 hESCs differentiated *in vitro* using the EB method. For Taqman analysis of differentiation (**Supplementary Fig. 9**), for a given EB-differentiated iPSC line, the expression of each gene is calculated relative to the expression of that gene in the undifferentiated sample for the same iPSC line. *GAPDH* was used for normalizing TaqMan data.

### *In vivo* pluripotency characterization

Teratoma formation in immunodeficient mice, harvest, and analysis was performed as previously described^45^. Briefly, iPSCs from 3 wells of a 6-well plate were collected with dispase in 50 μL of DMEM/F12 media. Before injection, 30 μL of Matrigel was added to the sample. Cells were injected subcutaneously in the rear flank of a NOD/SCID GAMMA C^-/-^ mouse. Teratomas were removed 6-12 weeks postinjection, fixed, sectioned, and evaluated histologically by hematoxylin and eosin staining.

### Northern blotting

iPSCs were treated with 100 μg/mL cycloheximide (Sigma C1988) or 1:1,000 dimethyl sulfoxide (DMSO) (Sigma D8418) for 3 hr prior to RNA collection. Total RNA was isolated from iPSCs using the RNeasy Mini Kit (Qiagen 74104) followed by mRNA collection using the PolyATtract mRNA Isolation System (Promega Z5310) according to the suppliers’ instructions. Northern blot analysis was performed using the NorthernMax-Gly kit (ThermoFisher AM1946). Two hundred ng of mRNA for each condition was run on a 1% glyoxyl/DMSO agarose gel and transferred onto a nylon membrane. Blots were probed with a *SLC9A6* or *GAPDH* probe labeled with α-^32^P UTP (Perkin Elmer) by T7 *in vitro* transcription of PCR-amplified cDNA. *SLC9A6* PCR primers for probe generation were forward: 5’- caggctttgtcgtcagaagtt −3’ and reverse: 5’- taatacgactcactatagggagagccacgaaagggtacaacat - 3’. *GAPDH* PCR primers were forward: 5’- ctgagaacgggaagcttgtc −3’ and reverse: 5’- taatacgactcactatagggagaggtgctaagcagttggtgg −3’. Radioactivity was measured using phosphorimaging (Molecular Dynamics) and quantified in ImageJ.

### Splicing analysis

iPSCs were treated with 100 μg/mL cycloheximide (Sigma C1988) or 1:1,000 DMSO (Sigma D8418) for 3 hr prior to RNA collection. Total RNA was isolated from iPSCs using the RNeasy Mini Kit (Qiagen 74104) followed by reverse transcription using Superscript IV RT (ThermoFisher 18091050) according to the manufacturer’s instructions. cDNA was PCR amplified using the following primers for *SLC9A6*, forward: 5’- caatgggtgctgctactgga −3’ and reverse: 5’- attggcagctcttcccaagaa −3’. PCR products were resolved on a 2% agarose gel, purified using the QIAquick Gel Extraction Kit (Qiagen 74104), and Sanger sequenced through GENEWIZ.

### Immunoprecipitation and western blotting

iPSCs were lysed in buffer containing Tris, pH 7.5, 1% Triton X-100 (Sigma), 150 mM NaCl, 20 mM β-glycerophosphate, 2 mM NaVO_3_, and 1X Complete Protease Inhibitor Cocktail (Roche 4693116001). Alternatively, iPSCs were lysed in buffer containing: 50 mM Tris-HCl, pH 7.8, 137 mM NaCl, 1 mM NaF, 1 mM NaVO_3_, 1% Triton X-100, 0.2% Sarkosyl, 1 mM dithiothreitol (DTT), and 10% glycerol supplemented with protease inhibitor cocktail and phosphatase inhibitor. All cells were lysed for 30 min on ice and were centrifuged at 13,200 rpm for 15 min at 4°C to remove cell debris. Protein concentration was measured by BCA assay using the Pierce BCA Kit (Thermofisher 23225). For immunoprecipitation, 10 μg of custom-made rabbit anti-NHE6 antibody (C-terminal epitope: GDHELVIRGTRLVLPMDDSE, Covance 048)^15^ was conjugated per 1.5 mg Dynabeads M-270 Epoxy (Thermofisher 14301) overnight at 37°C. Alternatively, 4 μg of anti-NHE6 antibody was conjugated to 0.5 mg Dynabeads Protein G (Thermofisher 10003D) for 2 hr at room temperature. After washing the beads, 500 μg of protein lysate was incubated with beads for 2 hr at 4°C. Samples were washed with buffer containing 20 mM Tris, pH 7.4, and 120 mM NaCl and eluted with buffer containing 20 mM HEPES, pH 8, 9 M urea, 1 mM NaVO_3_, 2.5 mM Na_4_P_2_O_7_, and 1 mM β-glycerophosphate. Samples were run on a NuPAGE 4-12% Bis-Tris gel (Novex NP0321BOX) and transferred to a nitrocellulose membrane (Novex LC2000). Membranes were blocked in blocking buffer (LI-COR 927-4000), and proteins were detected with rabbit anti-NHE6 antibody (Covance 048)^15^ and mouse anti-α-tubulin (Sigma T6074). Western blots were analyzed using a LI-COR Odyssey Imaging System.

Sequential study of NHE6 on western blot after immunoprecipitation was conducted, as we determined that the antibody appears to cross-react with NHE7 in human cells. Immunoprecipitation preceding blotting substantially ameliorated this problem. Notably, the rabbit anti-NHE6 antibody is raised to a cytoplasmic tail epitope (GDHELVIRGTRLVLPMDDSE) that is downstream of the truncation mutants, and therefore, this antibody would not be expected to identify proteins that are truncated prior to the epitope. After extensive study of all in-house generated and commercial antibodies, the antibodies we used are the best available for our purposes. Attempts to generate an N-terminal antibody failed, and none of the commercial antibodies are raised to the N-terminus of NHE6.

### Homology modeling

NHE6 models were constructed using MODELLER-9v14 (ref. ^46^) based on two template structures, including the crystal structures of the *E. coli* Na^+^/H^+^ exchanger NhaA (PDB identifiers 1ZCD (ref. ^47^) and 4AU5 (ref. ^24^)). The alignment between NHE6 and NhaA was generated based on a previously published alignment between NhaA and NHE9, a close homolog of NHE6 (ref. ^25^). For each template structure, 100 models were built using MODELLER and assessed with Z-DOPE, a normalized atomic distance-dependent statistical potential based on known protein structures^48^. Models constructed based on 1ZCD had six segments that were variable in sequence and were thus excluded from modeling. They included the loops between TM1-2, TM3-4, TM5-6, TM6-7, and the N- and C-termini. This model covered 53% of the NHE6 sequence. Models built based on 4AU5 contained the loops between the TM, while the N- and C-termini were not modeled. This model covered 65% of NHE6 sequence. The models were visualized using PyMOL (Schrödinger, LLC). Finally, we performed conservation and hydrophobicity pattern analysis to further validate the model^25,49^.

### Residue conservation and hydrophobicity

The ConSurf server (http://consurf.tau.ac.il/) was used to calculate evolutionary conservation (default parameters)^50^. To allow for a more complete analysis of NHE6 conservation, a model that contains loops was used (4AU5 template). The computed conservation values were mapped onto the NHE6 model with UCSF Chimera^51^, using the scripts provided by ConSurf. The hydrophobicity profile was computed using Chimera with the Wolfenden hydrophobicity scale^52^. Hydrophobicity values were mapped onto the NHE6 model’s surface, and the residues were colored from orange (most hydrophobic), to white, to blue (most hydrophilic).

### Modeling of the NHE6-G383D mutation

The G383D sidechain was modeled with PyMOL on a fixed backbone of the NHE6 wild-type model from MODELLER. The mutated model was further refined with Molecular dynamics (MD) simulations, where short MD simulations in implicit membrane were used as a control to test whether significant sidechain rearrangements occur in our model. GROMACS4.5 (ref. 53) was run on the G383D models with varying orientations of the G383D aspartate rotamer as described previously^54,55^. Each model was subjected to a short 10 ns simulation with 501 conjugate gradient minimizations.

### Neuronal induction monolayer protocol

Human neurons were generated using the previously described dual SMAD inhibition monolayer protocol with some modifications^36^. Control and CS subclonal lines belonging to the same donor family were differentiated simultaneously. iPSC lines with less than 30 passages were used for all neuronal differentiation experiments. In brief, iPSC colonies were treated with Gentle Cell Dissociation Reagent (StemCell Technologies 07174) for 10 min at 37°C and 1 x 10^6^ cells were plated onto one well of a Matrigel-coated 12-well plate in Neural Maintenance Media (NMM) (1:1 DMEM/F-12 GlutaMAX (Gibco):Neurobasal (Gibco)/1X N2 supplement (Gibco 17502048)/1X B27 supplement (Gibco 17504044)/5 μg per mL insulin (Sigma I9278)/1 mM L-glutamine (Gibco 25030149)/500 μM sodium pyruvate (Sigma S8636)/100 μM nonessential amino acids solution (Gibco 11140076)/100 μM β-mercaptoethanol (Gibco 21985023)/50 U per mL penicillin-streptomycin (Gibco 15070063)) in the presence of 10 μM Y-27632 dihydrochloride (Tocris 1254). After one day, if the cells had reached confluency, the media was changed to NMM + 1 μM Dorsomorphin (Stemgent 04-0024) + 10 μM SB431542 (Stem Cell Technologies 72232). Cells were fed daily for 10 to 12 days, after which they were passaged using Gentle Cell Dissociation Reagent, plated at 1:1 in one well of a 6-well plate, and fed with NMM. Cultures were inspected daily for formation of neuronal rosettes and were treated with 20 ng per mL FGF2 (PeproTech 100-18B) for 4 days to expand neuronal stem cells. Neuronal cultures were expanded and cryopreserved at days 22, 26, 30, and 34 after neuronal induction.

### Immunofluorescence protocols for human iPSCs and neurons

For detection of pluripotency marker expression by immunocytochemistry, iPSCs were fixed directly in cell culture dishes with 3.7% paraformaldehyde for 10 min at room temperature and permeabilized for 10 min in permeabilization buffer (PBS containing 0.5% Triton X-100). Non-specific binding was blocked by incubation with blocking buffer (permeabilization buffer + 5% normal donkey serum) for 30 min, and cells were incubated overnight at 4°C with mouse anti-Oct4 IgG (Santa Cruz sc-5279; 1:250) and rabbit anti-Nanog IgG (Abcam ab21624; 1:100) primary antibodies diluted in blocking buffer. Following washing with PBS, cells were incubated for 30 min at room temperature with donkey anti-mouse-IgG-Alexa Fluor-488 (Life Technologies A21202) and donkey anti-rabbit-IgG-Alexa Fluor-546 (Life Technologies A10040) diluted 1:500 in blocking buffer containing 0.3 μM DAPI. Following washing with PBS, cells expressing OCT4 and NANOG were visualized by fluorescence microscopy on a Leica DM IRB inverted fluorescence microscope. For detection of neuronal markers, iPSC-derived neurons were grown on 8-well glass chamber slides (Nunc Lab-Tek 154534) or on autoclaved glass 12-mm round coverslips (Ted Pella 26023) coated with poly-L-ornithine (Sigma P4957) followed by laminin (Sigma L2020). iPSC-derived neuronal cultures were fixed with 4% paraformaldehyde at room temperature for 10 min, permeabilized with 0.25% Triton X-100 (Sigma) in PBS for 10 min, and blocked with 10% goat serum (Jackson laboratories)/0.01% Triton X-100/1X PBS for 1 hr. Primary antibodies were diluted in 2% goat serum/0.01% Triton X-100/1X PBS and incubated for either 2 hr at room temperature or overnight at 4°C. The primary antibodies used were: chicken anti-MAP2 (Millipore AB5622), rabbit anti-PAX6 (Abcam ab5790), rabbit anti-TBR1 (Abcam ab31940), mouse anti-CTIP2 (Abcam ab28448), mouse anti-TAU1 (gift of Dr. Peter Dans), rabbit anti-HA (Cell Signaling 3724S), mouse anti-Ki67 (BD Biosciences 550609), rabbit anti-NeuN (Abcam ab177487), rabbit anti-activated-Cas3 (Cell Signaling 9661S), and mouse anti-NeuN (Millipore MAB377). After three, 5 min washes with 1X PBS, Alexa Fluor-conjugated secondary antibodies (Invitrogen) were diluted 1:1000 in 2% goat serum/0.01%Triton X-100/1X PBS + 1:1000 Hoechst (Invitrogen H3569) and incubated for 1 hr at room temperature. After PBS washes, cells were mounted with Fluoromount-G (SouthernBiotech 0100-01) and imaged on a Zeiss LSM 710 confocal laser scanning microscope.

### NanoString characterization of neuronal cultures

For each CS and paired control line, total RNA was collected from 1) iPSCs and 2) iPSC-derived neurons at 38 days post-induction using the RNeasy Mini Kit (Qiagen 74104). RNA was tested for quality using a Bioanalyzer (Agilent), and all samples had an RNA integrity number (RIN) of 9 or above. RNA was analyzed using a custom-designed NanoString platform containing probes for 358 genes related to cell fate, neurodevelopment, endosome biology, autophagy, lysosome biology, neurodegeneration, Trk and IGF signaling, and NHE family members. For each sample, 100 ng RNA was hybridized to the customized capture probe and reporter probe library for 12 hr at 65°C. Target-probe hybrids were immobilized onto a streptavidin-coated slide and imaged using the nCounter Digital Analyzer. NanoString data were analyzed with nSolver software. First, RNA counts were normalized to internal control sequences to account for variation in hybridization between samples. The geometric mean of the positive controls within each lane was calculated. Next, the average of the geomeans across lanes was calculated. Dividing the average across lanes by the geomean within each lane gives a lane-specific scaling factor. Counts within each lane were multiplied by the lane-specific scaling factor. Second, to account for variation in sample input, RNA counts were normalized to a set of housekeeping genes, including: *B2M*, *GUSB*, *POLR2A*, *GAPDH*, *HPRT1*, *RPL13a*, and *RPL27*. A lane-specific scaling factor was then calculated as above, and counts within each lane were multiplied by the respective scaling factor.

### Hierarchical clustering

Gene expression was measured for the 358 genes in the custom-designed NanoString panel in iPSCs and iPSC-derived neurons. Gene expression was Z-score transformed and used to perform agglomerative unsupervised hierarchical clustering using the Euclidean distance metric and average linkage criteria. Clustering was visualized using Java Tree View.

### Weighted Gene Co-expression Network Analysis (WGCNA)

Network analysis was performed using the WGCNA package in R and signed networks^34^. The construction of a network was based on the scale-free network concept indicating that a limited number of genes in this network are highly connected^33^. The advantage of this scale-free network is its robustness with respect to the choice of power value β. The relationship of each gene with disease status (iPSC/CS trait) was calculated by the Pearson’s correlation. A soft-threshold power of 12 (R^2^ > 0.8) was used for the frequency distribution of network connectivity to achieve approximate scale-free topology. An adjacency matrix was built with power function on a similarity matrix represented by Pearson’s coefficient of correlation. The network dendrogram was created using average linkage hierarchical clustering on the topological overlap dissimilarity matrix. Modules were defined as branches of the dendrogram using the Hybrid Dynamic Tree Cut method and signed networks^34,35^ with a deep split parameter of 2, a merged tree height of 0.9, and a minimum module size of 30 genes. Each qualified tree branch formed a module, as shown by color bands in the heatmap (Fig. 4c). Genes that were not in a specific module were assigned the color gray. Modules were summarized by their first principal component (module eigengene). Module membership represents the connectivity of a gene within a module. The intramodular connectivity was calculated by Pearson’s coefficient of correlation of a gene expression profile and the eigengene of that module.

### Time-lapse imaging of neurite outgrowth dynamics

iPSC-derived neurons were dissociated using Gentle Cell Dissociation Reagent or Accutase (StemCell Technologies) 32 days after neuronal induction. Thirty thousand cells were plated on 15-mm coverslips containing 50 μm laminin stripes that were patterned using micro-contact printing as previously described^56^. Cells were cultured in NMM for 12 hr, at which point a half media change was performed. Cells were then imaged using phase contrast on an inverted Nikon Eclipse TE2000-S microscope equipped with a motorized stage and with an enclosed custom humidified chamber to maintain 37°C and 5% CO_2_. Volocity software version 6.3.0 was used to acquire time-lapse series using a Hamamatsu Orca-ER camera, Orbit shutter controller, and a Ludl stage controller. For each sample, 12 Z-stack planes were acquired at 4 different XY positions every 5 min for 12 hr. Positions were selected based on the number of cells in the plane. Planes with too many cells crowded together were not used, as it was important to be able to measure dynamics of individual neurites. Neurons in which the processes were outside of the field of view were not used either. Imaging was performed simultaneously for each CS/control pair. Neurite tracking analysis was performed using the neurite outgrowth plugin in ImageJ (http://www.imagescience.org/meijering/software/neuronj/). The growing ends of the neurites were tracked for multiple neurons for the first 2 to 5 hr of each time-lapse movie.

### NHE6 re-expression and neuronal arborization

iPSC-derived neurons grown to 32 days after neuronal induction were transfected with a plasmid encoding for GFP alone or together with a plasmid encoding for human NHE6 (NM_006359) tagged C-terminally with HA (pReceiver-M07 mammalian expression vector) using Lipofectamine 3000 (ThermoFisher L30000001). Five days after transfection, cells were fixed and stained with rabbit anti-HA (Cell Signaling 3724S). Z-stack images were acquired using a Zeiss LSM 710 confocal laser-scanning microscope and traced for morphological analysis and Sholl analysis using Neurolucida software (MBF Bioscience).

### Endosomal acidification analysis in iPSCs

To measure changes in endosomal pH, iPSCs were loaded with FITC-conjugated transferrin (FITC-Tf) (ThermoFisher T2871) and Alexa Fluor-546- conjugated transferrin (Alexa-546-Tf) (ThermoFisher T23364) as previously described^15,32^. Briefly, cells were incubated with 66 μg/mL FITC-Tf and 33 μg/mL Alexa-546-Tf for 10 min, washed with PBS, collected using Gentle Cell Dissociation Reagent, and resuspended in PBS on ice. A standard curve was generated by resuspending cells incubated with FITC-Tf and Alexa-546-Tf for 30 min in standard buffer solutions containing: 125 mM KCl, 25 mM NaCl, 10 μM monensin, and either 25 mM HEPES (for standards pH 7.5 and 7.0) or 25 mM MES (for standards pH 6.5, 6.0, 5.5, and 5.0) and adjusted to a final pH using 1 N NaOH or 1 N HCl. The mean fluorescence intensity of FITC-Tf and Alexa-546-Tf was measured by flow cytometry using a BD FACS Aria III as previously described^32^. FlowJo software was used to gate on the live population of cells and calculate the mean fluorescence intensity for FITC and Alexa-546. The ratio of the mean fluorescence intensity of FITC to Alexa-546 was calculated for each sample, and endosomal pH was calculated using the standard curve.

### Sequencing of patient-derived and genome-edited iPSC lines

Genomic DNA was purified from 5 x 10^3^ iPSCs collected using Gentle Dissociation Reagent and washed one time with PBS before adding 30 μl of DNA extraction solution (Epicentre QE09050). Alternatively, DNA was extracted using Qiagen genomic-tips (Qiagen 10223). Primers for PCR amplification of the different exons of *SLC9A6* have been previously described^2^ and were exon 2: forward 5’- atccatagttatgcgtgggg −3’ and reverse 5’- ctcctggatcattttgctgc −3’; exon 3: forward 5’- tggaatatgaggtcttctaggga -3’ and reverse 5’- aggagaggagaaaggaggt -3’; exon 9: forward 5’- tggtggccttgcttaaactg -3’ and reverse 5’- ccaaaatgagtagatagcatcttgaac -3’; exon 11: forward 5’- tgagttccctaaaatagtctttcacc -3’ and reverse 5’- ttacatcaagctggcacacg -3’; exon 12: forward 5’- agaacgtgctttcttgctctg -3’ and reverse 5’- tggaaagatatccctcaaagtc -3’; exon 14: forward 5’- gcctctatttctgattctccaaag -3’ and reverse 5’- caaaacacttgttgttccattaaactg -3’. PCR amplicons were Sanger sequenced through GENEWIZ.

### Genome editing of human iPSCs to correct a mutation in *SLC9A6*

CRISPR/Cas9-based technology was used to introduce the wild-type *SLC9A6* sequence to an iPSC line derived from an individual harboring a c.1414dupA/p.R472fsX4 mutation in exon 11 of the *SLC9A6* gene. The guide RNA (gRNA) was designed according to the web tool (http://www.genome-engineering.org). Oligos (5’- caccgtccctcttacttaatt -3’ and 5’- aaacaattaagtaagagggac -3’) for gRNA expression were cloned into the pX458 plasmid allowing for co-expression of SpCas9-2A-GFP (Addgene 48138). The editing activity of the plasmid was validated in HEK293T cells by T7E1 assay. A donor ssODN (5’- cattgtgaatggttgcttgaagtgggcaagagaagaaagttacaacttaccagcaaacatcatcatgtgttgaaaatttgatccaatcttactccttcttcctaggtttagtaagagggacaaggggtaaatattggcagctcttc -3’) was designed to contain the desired wild-type *SLC9A6* sequence, flanked with homology arms to the targeted genomic region. The ssODN was also designed to contain silent mutations to prevent re-targeting by Cas9 and an *Avr*II restriction site to facilitate identification of targeted clones. For gene editing, a single-cell suspension of iPSCs was prepared using Accutase and 1 x 10^6^ cells were nucleofected with 2.5 μg of the plasmid and 2.5 μg of donor ssODN using program CA137. Forty-eight hr postnucleofection, GFP-positive cells were isolated by FACS and replated at cloning density in hESC media containing 20% KOSR, 4 ng/mL bFGF, and 10 μM Y27632 in 6-well dishes coated with 0.1% gelatin containing 187,500/well mitomycin C-inactivated CF1 MEFs. After 1-2 weeks, single clones with stereotypical iPSC morphology were manually excised and transferred to mTeSR1/Matrigel culture conditions for genotyping, expansion, and cryopreservation. Correctly targeted clones were identified by PCR, enzyme digestion, and Sanger sequencing.

### Generation of *SLC9A6* mutations in control iPSCs using CRISPR/Cas9 genome-editing technique

Human iPSCs derived from fibroblasts of a healthy male (Applied StemCell, Cat#: ASE-9203) were cultured and electroporated with a DNA mixture of plasmids allowing for expression of SLC9A6.g3,g11 and the Cas9 gene. Cells were selected by puromycin for a period of 24 to 48 hr two days post transfection. A small portion of the cell culture, presumably with mixed population, was subjected to genotype analysis. Once the mixed culture showed indels, it was subjected to a colony cloning process by picking and seeding each visible colony into a well of a 96-well plate. The cells were allowed to grow for 7 to 10 days or until a colony of workable size was formed. A fraction of cells from each colony was subjected to genotype analysis. Positive clones were expanded, and a portion of the cells was submitted to sequencing to confirm the desired genotype. Briefly, genotyping was performed as follows. Genomic DNA from single-cell colonies was isolated and used to amplify a 416-base pair DNA fragment using primers: forward 5’- ctgcatttcatctaaaaatcagttctgagctacag -3’ and reverse 5’- ccattacttattttctcctggatcattttgctgc -3’. PCR products were resolved on agarose gels and subjected to Sanger sequencing to screen for deletion mutants. The raw PCR products of the clones giving single bands were subjected to Sanger sequencing, and the results showed that there were various types of deletions. Clones that gave poor sequencing readouts or had deletions that did not result in frameshifts or truncations were eliminated from further analysis. The remaining clones showing frameshifts and premature stop codons resulting in short peptides were presumably knockout clones. To identify isogenic wild-type control clones, select clones that may not have been modified in the gene-editing process were picked for Sanger sequencing. Sequencing results indicated that two clones showed the identical sequence as the parental cells in the target region; cells from these clones were used as controls.

### Generation of *SLC9A6* knockout HEK293T cell lines using CRISPR/Cas9-based genome editing

Applied StemCell generated control and SLC9A6 knockout HEK293T cell lines. Briefly, the gRNA construct YH58.NHE6.g2 (5’- agaggctagagccatgg↓acgagg -3’; the arrow indicates the cleaving site) was used. This construct showed 57.2% cleavage activity and was transiently transfected into HEK293T cells, together with a plasmid encoding for Cas9. Cells were transiently selected with 2 μg/mL puromycin. Single-cell colonies were screened by genotyping. Genomic DNA from single-cell colonies was isolated and served as a source for amplification of a 477-bp DNA fragment using primers hSLC9A6_1F 5’- cctttcccgtgagccctcg -3’ and hSLC9A6_2R 5’- gctgcttcgcctcctctgc -3’. PCR products were then sequenced. Among 43 single-cell clones, clone 31 was identified as a homozygous hSLC9A6 knockout, with one ‘A’ addition at both or more alleles at the g2 cutting site; clone 35 was identified as a heterozygous hSLC9A6 knockout, with G/GACG/GACGAGGA deletion (−1/−4/−8) at each allele at the g2 cutting site; and clone 38 was identified as a clone with no mutations, thus providing an isogenic control clonal line (**Supplementary Fig. 15**).

### Generation of constructs encoding for the NHE6-G383D missense mutation

Constructs encoding for NHE6-G383D-HA, NHE6-G383D-GFP, and NHE6-G383D-mCherry were generated by PCR-mediated mutagenesis using the Quickchange II Site-Directed Mutagenesis Kit and according to the manufacturer’s instructions. Constructs were verified by Sanger DNA sequencing. Primers used for mutagenesis were as follows: hNHE6-g1148a_R 5’- acaatactgcaactacatctgtgaagccccatgct -3 and hNHE6-g1148a_F 5’- agcatggggcttcacagatgtagttgcagtattgt -3’.

### Dimerization assay for NHE6-WT and NHE6-G383D

HEK293T cells were transiently transfected using Lipofectamine 2000 with the various expression constructs encoding for GFP- or HA-tagged NHE6 as indicated in Fig. 6. After 24 hr, cells were harvested and then lysed in immunoprecipitation lysis buffer (50 mM Tris-HCl, pH 7.8, 137 mM NaCl, 1 mM NaF, 1 mM NaVO_3_, 1% Triton X-100, 0.2% Sarkosyl, 1 mM DTT, and 10% glycerol) supplemented with protease inhibitor cocktail and phosphatase inhibitor for 30 min on ice. Cell debris was pelleted by centrifuging the cell lysates at 13,200 rpm for 15 min at 4°C. The clarified cell lysates were then incubated with 15 μL (0.15 mg) of Pierce Anti-HA Magnetic Beads overnight at 4°C. The Anti-HA Magnetic Beads were then washed three times with 1X TBST (0.05%), boiled in Bio-Rad 2X sample buffer at 95°C for 5 min, and loaded onto 4-12% SDS-PAGE gels. Ten percent of the whole-cell lysate was loaded for input detection. Following electrophoresis, gels were transferred to nitrocellulose membrane for western blotting. Membranes were probed using antibodies against GFP and HA and imaged using the Li-CoR Odyssey Imaging System.

### Stability assay for NHE6-WT and NHE6-G383D protein

To investigate whether the G383D mutation altered NHE6 stability, a cycloheximide-based assay was used to measure the half-life of NHE6-WT vs. NHE6-G383D protein. Briefly, HEK293T cells were transiently transfected using Lipofectamine 2000 with constructs encoding for NHE6-WT-HA or NHE6-G383D-HA. Twenty-four hr post-transfection, cells were treated with 100 μg/mL cycloheximide for 0, 2, 4, or 8 hr. HEK293T cells were then harvested and processed for western blot analysis as described above for the dimerization assays.

### Endosomal pH rescue assays in HEK293T cells

The HEK293T control cell line (clone 38) and homologous hSLC9A6 knockout cell line (clone 31) were used to measure the effect of the NHE6-G383D mutant on endosomal pH. HEK293T hSLC9A6 knockout cells were transfected with construct(s) encoding for mCherry-tagged NHE6 as follows: NHE6-WT-mCherry alone, NHE6-G383D-mCherry alone, or NHE6-WT-mCherry and NHE6-G383D-mCherry together (construct ratio of 1:2). Cells were incubated for 24-36 hr following transfection, after which they, as well as control cells and untransfected hSLC9A6 knockout cells, were analyzed for endosomal pH using a flow cytometry-based assay as described above under “Endosomal acidification analysis in iPSCs.” FlowJo software was used to gate on the live population of mCherry-positive cells and calculate the mean fluorescence intensity for FITC and Alexa-546. The ratio of the mean fluorescence intensity of FITC to Alexa-546 was calculated for each sample, and endosomal pH was calculated using the standard curve.

### Microscopy-based analysis of endosomal pH in iPSC-derived neurons

To analyze endosomal pH in iPSC-derived neurons using a microscopy-based approach^32^, iPSC-derived neurons were first plated on glass-bottom imaging dishes (Nunc) and incubated in NMM without B27 and N2 supplements for 30 min. The media was then replaced with complete NMM containing 66 μg/mL FITC-Tf (ThermoFisher) and 33 μg/mL Alexa-546-Tf (ThermoFisher). After 10 min, the cells were washed 3 times with warm PBS, and the media was replaced with NMM without phenol red. Cells were imaged live using a Zeiss LSM710 confocal laser-scanning microscope. To calculate a standard curve, control iPSC-derived neurons were incubated with FITC-Tf and Alexa-546-Tf, and imaged in buffers of pH 5.0, 5.5, 6.0, 6.5, 7.0, and 7.5 (see above under “Endosomal acidification analysis in iPSCs”). The fluorescence intensity of each fluorophore was measured within endosomes using ImageJ. Endosomal pH was determined by calculating the ratio of FITC fluorescence intensity to Alexa-546 fluorescence intensity and extrapolating to the standard curve.

### Neuronal arborization and growth factor treatment

iPSC-derived neurons were plated on coverslips at 40,000 cells/cm^2^ and cultured in NMM. After 24 hr, media was replaced with NMM containing vehicle, IGF-1 (20 ng/ml) (PeproTech 100-11), or BDNF (10 ng/ml) (PeproTech 450-02). A half change of the respective media was performed the following day, and cells were fixed for staining 72 hr after vehicle or growth factor treatment. Immunocytochemistry and microscopy were performed as described above using chicken anti-MAP2 (Millipore AB-15452) and mouse anti-TAU1 (gift of Dr. Peter Dans). Neurite length and branching were quantified by tracing imaged cells using Neurolucida software.

### Statistics

NanoString gene expression differences were tested for significance with a two-tailed Welch’s t-test followed by Benjamini-Yekutieli FDR correction. Fold changes with an adjusted *P*-value < 0.05 were considered significant. Analysis of Sholl data was performed using a two-way ANOVA with Bonferroni multiple comparisons test. Neurite length and branching were compared using two-tailed Welch’s t-tests.

## Data availability

The NanoString data are included in this published article (and the accompanying Supplementary Information files). Additional datasets regarding endosomal pH assays and neuronal arborization and neurite growth studies are available from the corresponding author upon request.

## Acknowledgements

We thank the families who participated in this study. The study was supported by the following: Aspire I Junior Faculty Award, Center of Biomedical Excellence Dietary Supplements and Inflammation-NIGMS P20GM103641 Pilot Project Award, and SC INBRE NIGMS P20GM103499 Pilot Award (S.B.L.); NIGMS Postbaccalaureate Research Education Program (PREP) R25GM064118 (S.A.); Brown Institute for Brain Science Pilot Award and Brown University Seed Award (D.H.-K.); NIGMS R01GM108911 (A.S.); and Burroughs Wellcome Fund Career Award for Medical Scientists 1006815.01, NIMH R01MH105442, NIMH R01MH102418, NIMH R21MH115392, and Angelman Syndrome Foundation General Research Grant (E.M.M.). This study was also supported by the Hassenfeld Child Health Innovation Institute at Brown University, the Brown University Flow Cytometry and Sorting Facility, and the Cincinnati Children’s Hospital Medical Center Pluripotent Stem Cell Facility.

## Author Contributions

S.B.L., A.M.M., and E.M.M. designed the study. S.B.L., A.M.M., L.M., and L.I.v.D. performed the majority of the experiments and analyses. Q.W. and E.D.G.U. contributed to the bioinformatics-based analysis of NanoString data. L.L.L. and D.H.-K. contributed to the time-lapse imaging. L.M. and M.S. contributed to characterization of the iPSC lines, in particular, the genome-edited lines. L.M. performed studies relating to analysis of HEK293T cells. D.N., M.H.C., and P.B.-V. contributed to analysis of data relating to the iPSC lines. M.F.P. contributed to the recruitment of patients and obtaining samples for generating the iPSC lines. R.N.J. conducted statistical analyses. S.A. and A.S. conducted the structural modeling. S.B.L., A.M.M., and E.M.M. wrote the manuscript. All authors reviewed and approved the manuscript.

## Competing Interests

The authors declare no competing interests.

